# Microbial communities network structure across strong environmental gradients: How do they compare to macroorganisms?

**DOI:** 10.1101/2021.06.08.445284

**Authors:** C.M Arboleda-Baena, M. A Freilich, C.B Pareja, R Logares, R De la Iglesia, S.A Navarrete

## Abstract

The way strong environmental gradients shape multispecific assemblages has allowed us to examine a suite of ecological and evolutionary hypotheses about structure, regulation, and community responses to fluctuating environments. But whether the highly diverse co-occurring, free-living microorganisms are shaped in similar ways as macroscopic organisms, across the same gradients, has yet to be addressed in most ecosystems. The ‘everything is everywhere’ hypothesis suggests they are not, at least not to the same extent. Here we characterize the structure of intertidal microbial biofilm communities and compare the intensity of zonation at the ‘species’ level, changes in taxonomic diversity and composition at the community level, and network attributes, with those observed in co-occurring macroalgae and invertebrates. At the level of species and OTUs, for dominant macro and microorganisms respectively, microbes showed less variability across the tidal gradient than macroorganisms. At the community-level, however, microbes and macro-organisms showed similarly strong patterns of tidal zonation, with major changes in composition and relative abundances across tides. Moreover, the proportion of ‘environmental specialists’ in different tidal zones was remarkably similar in micro and macroscopic communities, and taxonomic richness and diversity followed similar trends, with lower values in the high intertidal zone. Network analyses showed similar connectivity and transitivity, despite the large differences in absolute richness between the groups. A high proportion of positive co-occurrences within all tidal zones and mostly negative links between the high and low tidal zones were observed among habitat specialist taxa of micro-and macro-organisms. Thus, our results provide partial support to the idea that microbes are less affected by environmental variability than macroscopic counterparts. At the species-level, the most common microbe species exhibit less variation across tides than most common macroscopic organisms, suggesting the former perceive a more homogeneous environment and/or are more resistant to the associated stress. At the community-level, most indicators of community and network structure across the gradient are similar between microbes and macro-organisms, suggesting that despite orders of magnitude differences in richness and size, these two systems respond to stress gradients, giving rise to zonation patterns.

## Introduction

Understanding the mechanisms and processes responsible for patterns of abundance and distribution of species across environmental gradients is one of the main goals of community ecology (Mittelbach and McGill 2019), and one of the most striking patterns is the zonation of dominant organisms that occupy different parts of the environmental gradient. These patterns have served as the basis to test core hypotheses about the organization of communities (e.g. Whittaker 1959) and the ecological processes shaping fundamental and realized species’ niches, such as physiological tolerances and species interactions (Connell 1961a, 1961b, Menge and Sutherland 1987). However, this deep understanding of communities of macroscopic organisms contrasts sharply with the still scarce knowledge about communities of free-living microorganisms (Fontaneto 2011, Hanson et al. 2012). The ‘everything is everywhere, but the environment selects’ hypothesis, set forth by Gerhard Baas Becking in the early 1930s, has inspired much of the microbial biogeographical and ecological research of the past decades (de Wit and Bouvier 2006, O’Malley 2007) and suggests that microbial communities are primarily determined by moderate rates of dispersal and secondarily by environmental variation (selection). It is clear that microbial biodiversity is shaped by environmental gradients, especially by those factors that alter soil or water pH (Rousk et al. 2010, Krause et al. 2012), temperature (Logares et al. 2020), and resource availability (Follows et al. 2007). The local environment ‘selects’ which microorganisms are present and, like in macroscopic organisms, patterns of abundance and composition across environmental gradients result from the constraints set by abiotic conditions and the interactions among co-occurring microbes (Fontaneto 2011, Hanson et al. 2012, Mandakovic et al. 2018). Yet, the question remains, are communities of microbial organisms less structured across the same environmental gradients than their co-occurring macroscopic counterparts? If both vary across sharp environmental gradients, are the patterns of community structure towards the stressful and benign ends of a gradient similar to those observed in macroscopic organisms? Addressing these and other similar questions allows us to challenge our ecological models and get valuable insight into how different is the organization between the micro and macroscopic worlds.

Probably the most studied zonation pattern is the one occurring between the highest and lowest tides along most marine rocky shores of the world (Barnes and Hughes 1999, Raffaelli and Hawkins 2012). In this habitat, environmental conditions range from entirely aquatic to completely terrestrial in a few meters (Thompson et al. 2002, Harley and Helmuth 2003); and, within their absolute tolerance limits, species become consistently more abundant at some tidal levels than others (Stephenson and Stephenson 1949, Connell 1961a, 1961b). Also, co-occurrence network analyses show modules (cluster of species highly connected) related to a specific zonation band due mainly driven by shared environmental preferences (Freilich et al. 2018). Beyond the tidal regime that sets the stage for the stress gradient (Denny and Paine 1998, Raffaelli and Hawkins 2012), intertidal zonation of macroscopic organisms results from multiple interactive physical factors, including wave exposure and the associated mechanical stresses, desiccation (Evans 1947), temperature (Wethey 1983), and solar radiation (Santelices 1990), to name a few. Additionally, species interactions, including interspecific competition for space (Connell 1961b), food acquisition (Underwood 1972), facilitation (Bertness and Leonard 1997), and predation (Connell 1961a, Paine 1966), profoundly modify intertidal zonation. As a result, zonation of organisms results from a combination of environmental stressors affecting differentially some species, and the propagation of these effects throughout the community via species interactions. To what extent the intertidal gradient imposes similar patterns of community structure on macro and microscopic communities has not been addressed.

Intertidal microbial communities on rocky shores can be found as individual cells or forming well-defined biofilms that can grow on virtually all surfaces (Callow and Callow 2011). Biofilms are composed of taxa belonging to the three domains of life, associated with extracellular polymeric substances (EPS), acquiring important emergent properties such as hydrophobicity, viscoelasticity, drug resistance among others (Davey and O’toole 2000, Schuster et al. 2019). Studies have shown that variability in abundance and the broad composition of intertidal epilithic biofilms are influenced by temperature, desiccation, UV radiation, waves (Thompson et al. 2004, 2005), and the concentration of ions and nutrients (Decho 2000, Dang and Lovell 2016). Competition and predation between microorganisms (Dang and Lovell 2016), competition with macroscopic sessile species (Callow and Callow 2011), and top-down control by grazing macroinvertebrates (Lubchenco 1978, Underwood 1984) all influence abundance and tidal distribution of major groups of microbial biofilms. However, most ecological studies have treated biofilms as homogeneous taxonomic units composed of few entities, precluding a community- or ecological-network approach comparable to macroscopic organisms. Since 2006, massive sequencing techniques allow us to explore and document the diversity of the microbial world as never before (Hug et al. 2016), opening the doors to address ecological questions. Here we compare patterns of zonation in taxonomic diversity, richness, and network structure of the microbial intertidal communities on wave-exposed rocky shores of central Chile and the well-documented patterns of co-occurring macroscopic organisms (Castilla 1981, Santelices 1990, Kéfi et al. 2015, Freilich et al. 2018). Using the previously defined zonation bands for macroorganisms, we test four hypotheses: 1) that across the same environmental gradient, microbial communities show proportionally less differentiation across tidal levels than their macroscopic counterpart, 2) that taxonomic richness and diversity do not follow similar trends across the gradient, 3) that the proportion of environmental specialist is higher in the macroscopic community than in microscopic organisms, 4) that co-occurrence networks of macroorganisms have modules corresponding roughly with zonation bands, while microbial networks do not form zonation-related compartments.

## Methods

The comparison of macro and micro-communities represents a large methodological, logistical, and economic (DNA sequencing) challenge. We propose and describe here a reproducible method to quantify microbial communities across tidal levels for comparison with macroscopic communities.

### Macroscopic community data

Quantitative comparisons with macroscopic communities were based on intertidal surveys conducted since 1998 across sites in central Chile from the 33° 42’ S, 71° 70’ W to 33° 59’ S, 71° 62’ W, including the study site for microscopic studies. Field methods have been described in detail (Broitman et al. 2001, Freilich et al. 2018). Briefly, surveys at each site quantified the density of mobile species and the cover of sessile species of all macroscopic organisms (>1mm) in 8-10 50 × 50 cm quadrats along transects parallel to shore at three tidal levels, low, mid, and high shore. Details of surveys included in this study are presented in **Appendix S1: Table S1**.

### Microbial community data

Microbial community studies were conducted in the wave-exposed rocky shore at Las Cruces, central Chile, 33° 50’ S, 71° 63’ W. The rock is formed by postmetamorphic granite with basaltic intrusions and high quartz content and tidal range is about 1.8 m, with semidiurnal regime. Since our goal was to compare the organization of microbial communities across different tidal levels, it was critical to set a similar successional time and reduce interference with surrounding macroscopic biota as much as possible. We therefore developed a simple method that can be replicated in future studies. First, we collected rocks from the same study site and cut them into coupons of 3 × 8 × 2 cm with a COCH Bridge saw machine that prevented overheating and potential mineral transformations. Rock coupons were washed with deionized water, dried, maintained at room temperature, and then attached individually, with a stainless-steel wire, inside stainless-steel cages 12 × 12 × 4 cm and 5 mm mesh. Cages prevented grazers and other macroscopic organisms from crawling on the rock coupons. Twelve rock coupons inside a separate stainless cage were affixed with a stainless-steel screw at each of three tidal zones haphazardly distributed along tens of meters parallel to the shore. Tidal zones followed the same levels used in macroscopic studies described above(Santelices 1990, Flores et al. 2019). Five rocks selected at random were not deployed in the field and were used as rock surface control. After 40 days of exposure, rock coupons were retrieved and, together with rock surface controls, were sonicated in sterile marine water, and the biofilm was recovered through 0.22 μm pore filters of hydrophilic polyethersulfone (Merck). Samples were preserved in liquid nitrogen at - 196°C for molecular analyses. A total of five cages with rock coupons were lost due to storms. A 40-day elapsed time has been shown to allow intertidal biofilm communities to reach a comparatively stable structure at the end of succession, and macroorganisms do not cover them yet (authors, personal observation).

### DNA extraction, 16S rRNA sequencing and molecular data analyses

DNA extraction from filters was conducted with the Phenol-Chloroform method (Fuhrman et al. 1988) and DNA samples were clean by the membrane dialysis method with a Millipore 0.025 uM filter (Devaney and Marino 2001). DNA concentration was measured with the Qubit HS dsDNA Assay kit in a Qubit 2.0 Fluorometer (Life Technologies, Carlsbad, CA, USA) and the DNA purity was quantified using a NanoDrop-1000 (NanoDrop Technologies, Wilmington, USA), according to manufacture protocols. The V4-V5 region of the 16S was amplified with the primers 515FB: GTGYCAGCMGCCGCGGTAA and 926R: CCGYCAATTYMTTTRAGTTT (Parada et al. 2016). Amplicons were sequenced in a MiSeq Illumina platform (2×300 bp). Both PCR and sequencing were done at the Dalhousie University CGEB-IMR (http://cgeb-imr.ca/). Sequence data were deposited at the European Nucleotide Archive (ENA) public database under accession number PRJEB45042.

### Macro and Micro community analyses

Amplicon reads were analyzed using the DADA2 pipeline (Callahan et al. 2016, Lee 2019), generating Amplicon Sequence Variants (ASVs) that were used as Operational Taxonomic Units (OTUs). Rarefaction curves (Simberloff 1978) for bacterial diversity were generated with the R package phyloseq v1.30.0 (McMurdie and Holmes 2013) and allowed us to define a sampling effort of 16,888 reads for microbial diversity comparisons across tidal levels (**Appendix S1: Table S2**). The number of different OTUs and macroscopic species per unit area (hereafter called ‘richness’), were separately compared among treatments (three tidal zones, fixed factor) with Welch ANOVA, and one-way ANOVA for macro and micro communities, respectively, since the former displayed heteroscedasticity. The Games-Howell and a Tukey’s posteriori tests were performed after significant ANOVA results, for macro and microbial communities, respectively. Species and OTUs Shannon diversity indices were compared among tidal zones with separate Welch ANOVAs and Games-Howell a posteriori tests for macro and microorganisms, as they both exhibited heteroscedasticity. To quantify tidal variation in abundance and thus the intensity of zonation patterns, we calculated coefficients of variation (CV) across tidal levels of the most abundant macro and microorganisms at the level of species and Orders. To characterize community compositional differences and general structure among tidal zones, we conducted multivariate ordinations on presence/absence data using Jaccard distance index, and on relativized species/OTU abundances using Bray Curtis distance for Non-Metric Multidimensional Scaling ordinations. To test for statistical significance in community composition and structure among the intertidal zones, we conducted permutational analysis of variance (PERMANOVA) (Anderson and Walsh 2013), followed by pairwise a posteriori tests using FDR correction (Benjamini and Hochberg (1995), separately for micro and macroscopic organisms. We also conducted quantitative Principal Coordinate Analyses (PCoA) to represent tidal zones in the multispecies space to recover the centroid that defined the compositional groups for macro and microorganisms. This allowed us to calculate multivariate Mahalanobis distances among community centroids at the three intertidal zones. All multivariate analyses were conducted with the R package vegan v2.5-6 (Oksanen et al. 2015).

### Co-occurrence Network analyses

To further compare the structure of macro- and microbial communities across tidal zones, we constructed co-occurrence networks for both groups using maximal information coefficient analysis applied to presence/absence data in the software MICtools (Albanese et al. 2018) to derive the strength and sign of correlation. Tidal zone specialists were identified with the Indicator Value (IndVal) (Dufrene and Legendre 1997) in the labdsv R package (Roberts and Roberts 2016), which identifies Species/OTUs associated to a given tidal zone based on fidelity and relative abundance. To compare network structures, we computed connectivity, the proportion of all possible edges that are realized in the network (Newman 2018), and transitivity, the ratio of the number of triangles to the paths of length two, which quantifies the clustering of the network (Wasserman and Faust 1994). Both metrics were calculated for micro and macroorganisms separately. We then identified the habitats specialists and not specialist in each network, according to the IndVal index, and represented positive and negative links within and between tidal zone compartments. To summarize patterns of positive and negative links, we calculated connectivity within and between entire compartments of habitat specialists and not specialist. The statistical significance of network connectivity and transitivity was evaluated against Erdos-Renyi random networks of the same size (Connor et al. 2017). The existence of compartments in the macro- and microbial networks was examined using the leading eigenvalue method (Newman 2006). Network metrics were computed using the R package iGraph (Csardi and Nepusz 2006).

## Results

### Zonation of dominant taxa

A total of 88 macroalgae (n= 40) and invertebrate (n = 48) species were found at the surveyed sites of central Chile (**Appendix S1: Table S3**). Dominant species in each zone were similar to those reported in previous studies (Castilla 1981, Broitman et al. 2001) (**Appendix S1: Table S3**). We obtained 472,864 good-quality sequences from 28 samples for microbial community analyses. After rarefying to 16,888 reads per sample, due to the size of the smallest dataset of one low intertidal zone sample, we obtained a total of 6,252 OTUs (**Appendix S1: Table S2**). The main phyla in the intertidal zone were Proteobacteria (Alphaproteobacteria and Gammaproteobacteria), Bacteroidetes, Cyanobacteria, and Planctomycetes (**Appendix S1: Figure S2, Table S3)**. Patterns of zonation, i.e., sharp changes in relative abundance, of the 14 most abundant macroalgae and invertebrate species across the tidal gradient were very apparent (**Fig. 1**). Although most species were present at all three tidal zones, they show high abundance at a single (e.g., *Ectocarpus silicosus*, *Nodolittorina peruviana*) or, at the most, two tidal zones (e.g., *Ralfsia californica, Siphonaria lessoni*) (**Fig. 1a**). Zonation was also apparent in the 14 most common OTUs, although the changes of abundance across tidal zones were attenuated compared to macroscopic organisms (**Fig. 1b**). This difference was reflected in larger coefficients of variation (CV) in macro-(mean= 123%) than micro-organisms (mean= 84%) (**Fig. 1c,d**). Pooling species and OTUs into the main Orders (**Appendix S1: Figure S1a,b)**still revealed patterns of specialization or zonation in macroorganisms, but only weakly so in microorganisms (mean CV: Macro= 125%, Micro= 62%) (**Appendix S1: Figure S1c,d).**A negative slope characterized the relationship between spatial variation in orders of macro- and micro-organisms and the number of species (S) within the Orders (**Fig. 2a,b**), but the relationship was statistically significant only for macroorganisms. The negative, buffering effect of diversity on variability across the gradient was similar in micro-and macro-organisms (similar slopes, ANCOVA interaction: p= 0.459), and a single linear fit across micro and macro-organisms was highly significant (CV= 150 – 66.8 + logS, R2= 0.43, p = 0.001, **Appendix S2: Figure S1)**, but this result should be taken with caution because of the inevitable collinearity produced by differences in richness (**Fig. 2c,d**). A detailed account of main macro-organisms and microbial orders and their variability across tidal zones is displayed in **Appendix S1: Figure S1.**

**Fig. 1.**
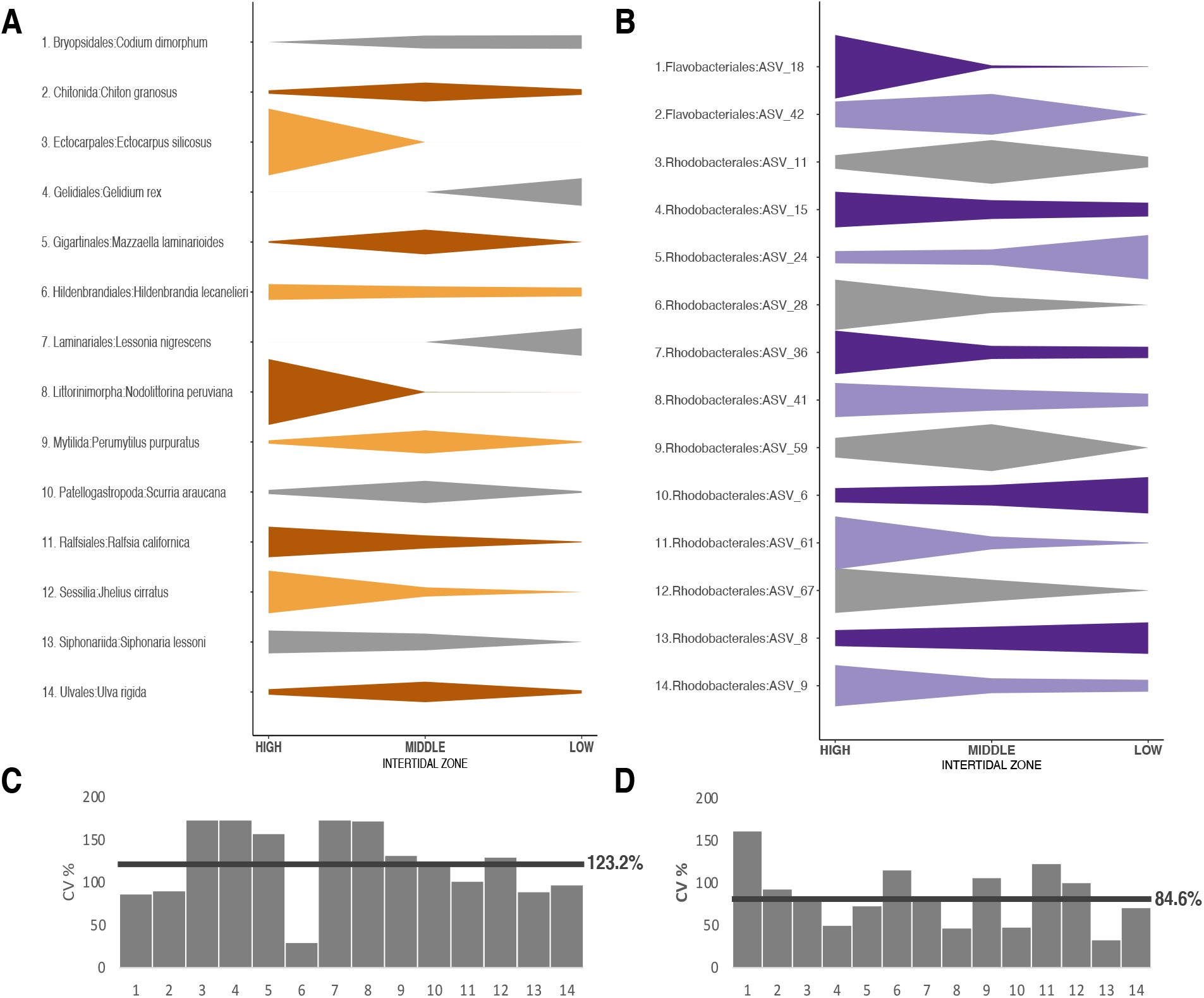
Zonation patterns of the 14 most abundant species (in terms of relativized cover, density, or reads of sessile invertebrates and macroalgal species, mobile invertebrate species, or OTUs, respectively), expressed here as relative to the maximum abundance observed in a given zone for **A)** Macroorganisms and **B)** Microorganisms. The coefficients of variation (CV%) across tidal zones for the same species and OTUs are shown for **C)** macro- and **D)** micro-organisms. The horizontal lines are mean CV for macroorganisms = 123.2% and microorganisms = 84.6%.

**Fig. 2.**
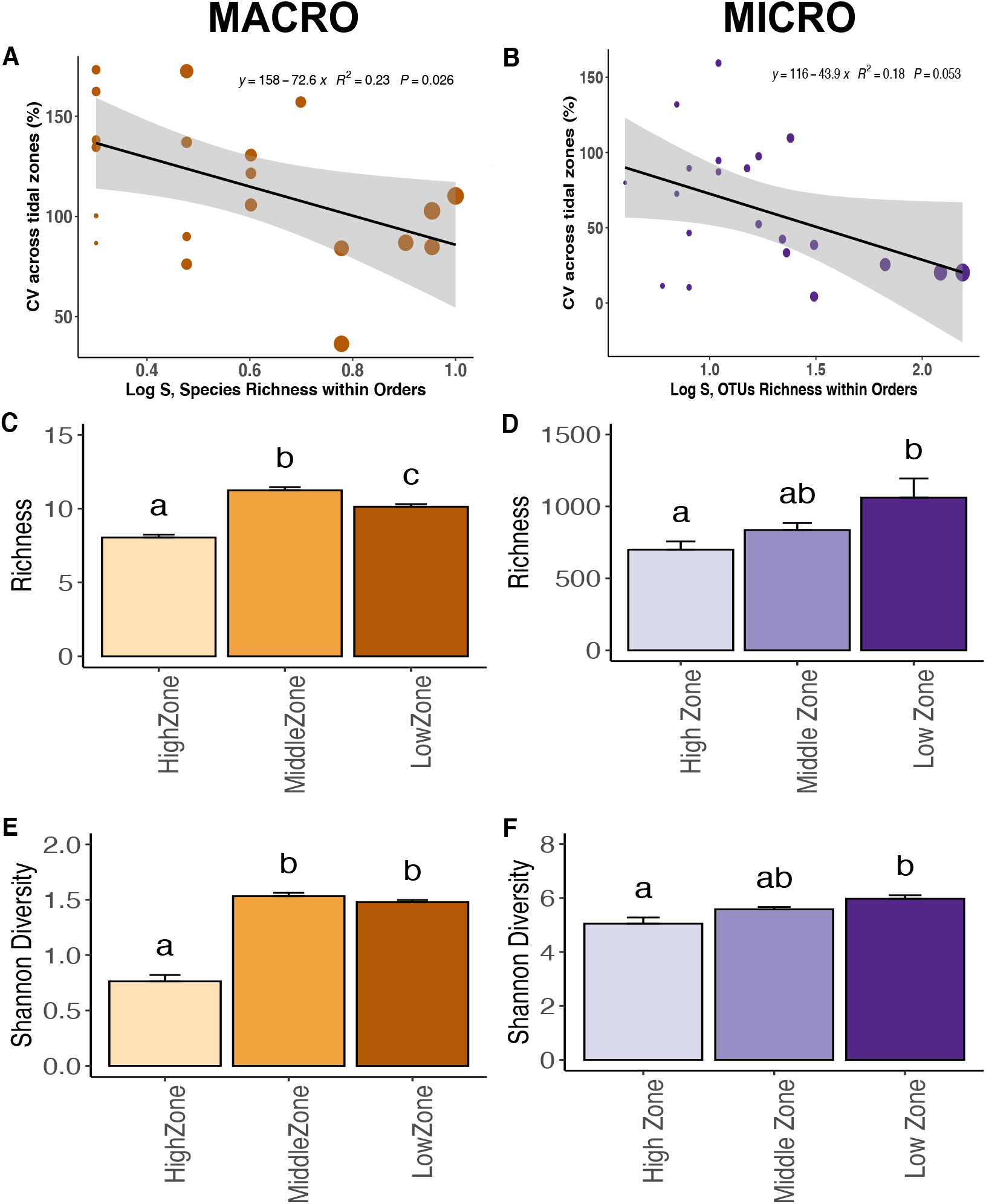
Linear regression between the coefficient of variation across tidal zones and the OTUs richness within Orders for macro- (**A**) and microorganisms (B). Mean (+ SE) richness (**C,D**) and Shannon diversity (**E,F**) of (**C,E**) macroscopic community and (**D,F**) microbial community in the high, middle, and low intertidal zones. Different letters above bars indicate significant differences with a posteriori tests (**C,E, F**: Games-Howell tests, **D**: Tukey test at the experiment wise error rate = 0.05).

### Diversity across tidal zones

Species richness of macroscopic communities was significantly different across tidal zones (**Fig. 2c**, Welch’s ANOVA, p<0.0001), with highest richness in the mid zone and lowest in the high zone. The richness of OTUs (using rarefaction of equal sampling effort) also showed significantly different values among intertidal zones (**Fig. 2d**, ANOVA, p=0.0209), with highest values in the low zone and lowest in the high zone. Microbial richness in the mid-zone reached intermediate levels and could not be statististical differentiated from values observed in the low or high zones. The Shannon diversity index for macroorganisms also showed significant differences among intertidal zones (**Fig. 2e**, Welch’s ANOVA, p<0.0001), with higher and similar values in the middle and low zones than the high zone (Games-Howell post hoc test, p=0.49). Microbial diversity also varied significantly among intertidal zones (**Fig. 2f,** Welch’s ANOVA, p=0.0143) with the highest values in the low zone and lowest in the high zone (Games-Howell tests). Intermediate diversity levels in the mid zone could not be statistically separated from other tidal zones by a posteriori tests. Richness and Shannon diversity were significantly different between microbial communities of the three intertidal zones and the rock surface control (**Appendix S2: Figure S2)**.

### Community and network structure

The composition and relative abundances of the macroorganisms community were different among the three tidal zones (**Fig. 3a,c**), with sharper separation between the low zone and the other two, and largest dispersion in the mid zone (**Fig. 3a,c**). These multispecic differences among zones were statistically significant (**Appendix S3: Table S1a,b**, PERMANOVA, p=0.0009 for Jaccard and Bray-Curtis distances), and paired comparisons showed that all intertidal zones differed from each other (FDR adjusted post-hoc tests: p=0.03 Jaccard distances and p=0.02 Bray-Curtis distances). The microbial communities tended to show similar patterns to their macroscopic counterparts, with a sharply defined low intertidal zone and increased overlap between the high and mid zones (**Fig. 3b,d**). In this case, the largest dispersion was observed in the high intertidal zone. Statistically significant differences were observed in microbial community composition and relative abundances (**Appendix S3: Table S1c,d**, PERMANOVA, p=0.0009 for Jaccard and Bray-Curtis distances), and paired comparisons also showed that all zones differed from each other (FDR adjusted post-hoc test, p=0.03 Jaccard and p=0.02 Bray-Curtis distances). Including the rock surface’s control in the analyses did not alter this general pattern **(Appendix S3: Figure S1).**

**Fig. 3.**
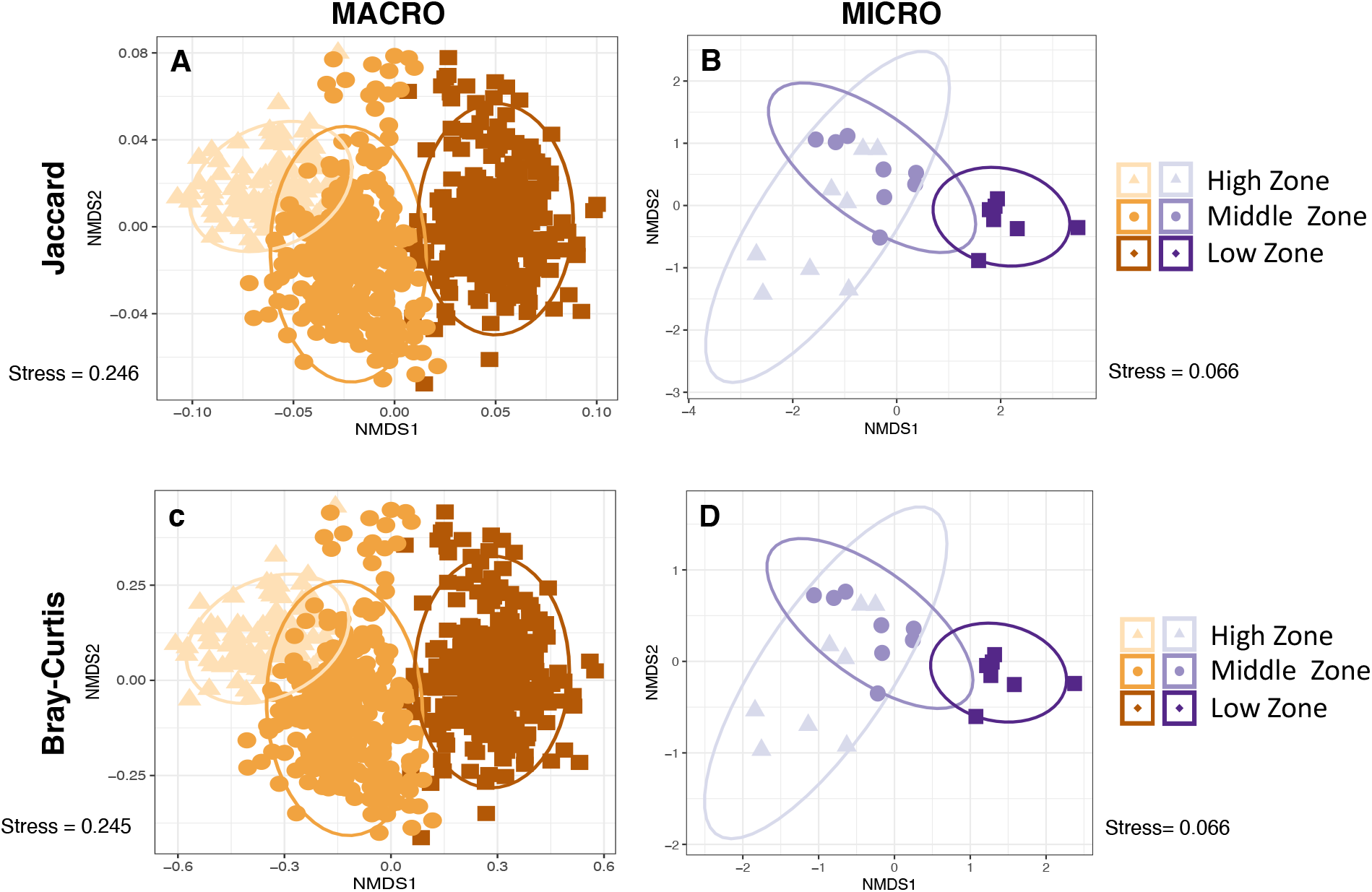
Compositional similarity of the different intertidal rocky shore zones for macroorganisms (N= 88 species) and microorganisms (N= 6,252 OTUs). Non-metric multidimensional scaling (NMDS) ordination plots based on presence/absence data using Jaccard distances (**A**, **C**), and on abundance data using Bray-Curtis distances (**B**, **D**). The symbols represent the different intertidal zones, surrounded by the ellipse of 95% confidence interval. Each observation is a survey unit (50 × 50 cm quadrats for macroscopic organisms, 3 × 8 cm rock surface coupons for microbes).

The connectivity and transitivity analyses of the co-occurrence networks showed similar connectivity, 0.216 and 0.222 for macro and micro-organisms, respectively, and the same undirected transitivities of 0.56. These transitivity values were in both cases significantly larger than the transitivity of random networks (p < 0.001). While most species and OTUs changed in abundance across tidal zones, our IndVal index analyses allowed us to classify them into habitat specialists or not specialist and identify these types of species/OTUs in the networks (**Appendix S4: Table S1a,b**). The fraction of habitat specialists in the high, mid, and low zones was remarkably similar between micro- and macroorganisms (**Fig. 4a,b**), with the highest number found in the low shore. In micro and macroorganisms networks, positive correlations dominated the links detected between taxa within all tidal zone compartments, while predominantly negative links were detected between species in the high and low shore compartments (**Fig. 4a,b**). In both networks, not specialist were positively and negatively connected with habitat specialists and other not specialist. These visual patterns were well captured in the connectivity summary within and between compartments. This showed high and generally positive connectivity within compartments, but strong and, on average, negative connection between the high and low zones (**Fig. 4c,d**). Macroorganisms restricted to the high intertidal and microorganisms restricted to the low intertidal are particularly highly connected within their groups of habitat specialists with connectivity of 0.73 and 0.58, respectively, and strong positive correlation (**Fig. 4c,d**). In both networks, the average correlation strengths between specialists and nonspecialists was very weak, although the values were higher in macro-organisms than in the microbial network (**Fig. 4c,d**), a pattern that was apparent in the visual representation of links. Despite these remarkable similarities, the two networks had different degree distributions, i.e., how the number of links of each node are distributed, with binomial distribution in macroscopic organisms and power-law degree distribution in microorganisms (**Appendix S4: Figure S1**). Modularity analyses of the entire networks showed a weak and non-significant relationship of compartments to the tidal zonation in both networks (**Appendix S4: Figure S2)**.

**Fig. 4.**
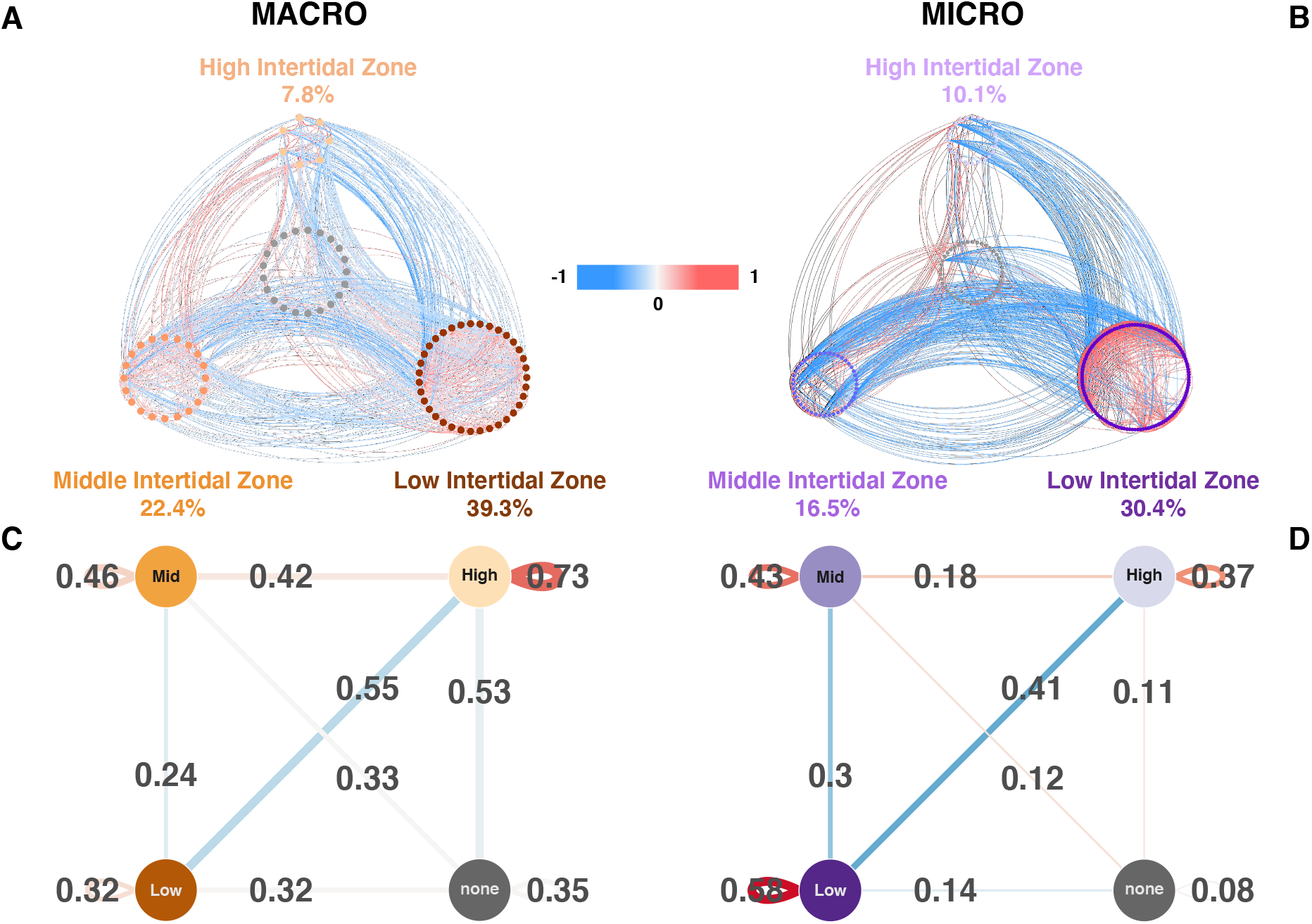
Co-occurrence networks of A) macroorganisms and B) microorganisms communities. The node color denotes the intertidal zone specialists at the 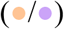 high, 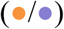 middle and 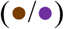 low intertidal zone. The gray nodes 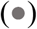 are species/OTUs without specialisms. The edge color denotes 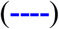 positive correlation and 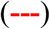 negative correlation. The magnitude of the correlation is shown by the intensity of the shade, as depicted in the legend between the two networks. Connectivity of subnetworks of C) macroorganisms and D) microorganisms communities. The edge width is proportional to the connectivity within and between each subnetworks of tidal height specialists. The edges are labeled with the connectivity. “None” indicates non-specialists. The edge color is proportional to the average correlation in subnetworks within and between tidal height specialists and non-specialists.

## Discussion

Our combination of intertidal surveys and experiments allowed us to characterize microbial communities and make sensible comparisons with macroscopic organisms across one of the best studied environmental gradients in the world. While it is true that these comparisons are constrained by methodological issues, which bound our conclusions, they still shed light on the processes driving ecological organization in microbial organisms and how different they may be from those affecting co-occurring macroscopic components of these communities. Our results partially support the idea that microbes are less affected by environmental variability than macroscopic counterparts. First, while a similar fraction of all micro-and macro-organisms can be considered as ‘habitat specialists’, i.e., largely restricted to a given tidal zone, the 14 most common microbial species exhibit much less variation across tides than the most common macroscopic organisms, suggesting the former perceive a more homogeneous environment and/or are more resistant to the associated stress. At the community-level, however, most indicators of community structure and attributes of co-occurrence networks across the gradient were remarkably similar between microbes and macro-organisms, suggesting that despite orders of magnitude differences in richness and size, these two systems respond to stress gradients at a multispecific level in similar ways. We discuss these results below but start with a brief discussion of the methods used, which must be borne in mind when drawing conclusions. Methodological approximation to intertidal biofilm studies has followed two general approaches. Firsts, removal of sections of natural, existing rocks from the environment with different exposure time (De la Iglesia et al. 2012, Taylor et al. 2014, Tan et al. 2015, Kerfahi et al. 2020), or the installation of artificial surfaces such as plexiglass, polystyrene, or glass slides (Zhang et al. 2013, Sanli et al. 2015). The first approach is useful for examining the composition of microbial communities but makes it difficult to make sensible comparisons across gradients because the successional time is unknown. The second approach allows for comparative studies, but the specificity of microbial communities for different surfaces limits interpretation to natural systems. Our study attempts to solve these issues using the same natural rock surface, with the same area, physical attributes, chemical composition, and the same exposure time for all treatments, allowing us to compare among treatments and extrapolate to natural rock surfaces. Still, some variation in micro-flows concerning a flat rocky platform is inevitable. Moreover, we opted to excluded macro-organisms and, in this manner, reduce interactions with macroscopic organisms, especially molluscan grazers. This was a difficult decision because it introduced the potential artifact of having a stainless-steel mesh reducing the light incidence and probably ameliorating stress on experimental rocks. We could devise no appropriate control for this artifact without a prohibitively large number of treatments. Future studies should therefore consider these potential cage effects and experimentally assess the effect of macroscopic organisms on reported patterns of microbial zonation. Finally, we decided to use large numbers of surveys for macroscopic organisms at the biofilm study site and nearby sites (see **Appendix S1: Table S1**), to have a larger sample for co-occurrence network analyses. This can be easily changed by considering only one or more sites and data are made available. Our broad conclusion is not altered by this consideration.

Our results indicated that dominant microbial organisms across the same environmental gradient showed comparatively less differentiation across tidal levels than their macroscopic counterpart. i.e., the tidal gradient strongly shaped the most abundant species and OTUs. The common explanation for this might be that the environmental conditions in the intertidal rocky shore perceived at the microscopic level are less stringent than those perceived by macroscopic organisms. While this is entirely possible, it is also possible that these organisms are more resistant to similar gradients in stress, which can only be assessed by experimentally manipulating stress levels. To what extent the cage used to exclude macroscopic organisms reduced the amplitude of the stress gradient must also be further investigated. Regardless of the mechanism accounting for differences in stress-related responses at the ‘species’ level, we observed a strong buffering effect of species richness within higher taxa, such as Orders, on the response to the stress gradient when considering micro and macro-organisms together. (**Appendix S1: Figure S1**), and that the effect was apparently homogeneous between these groups (non-significantly different slopes). Thus, there is an indication that in both groups, increased species redundancy confers resistance to environmental stress as a taxon. This is an interesting result that deserves further studies as it suggests similarities between micro and macroscopic worlds. However, it must still be taken with caution because the linear fit relationship was marginally non-significant for the microscopic organism when analyzed separately. These results also call for caution when analyzing microbial communities across space without resolving to OTUs level because aggregation at higher taxa is bound to support the everything is everywhere paradigm equivocally.

At the community-level, stress gradient responses of micro and macroscopic communities become more similar. The time exposure to radiation, high temperature, ultraviolet light, and desiccation are factors that could explain the similar trends across the gradient between these two communities because, in both group lowest values of abundance, richness and diversity were observed in the high intertidal zone, increasing to the middle and low tidal levels. Other studies supported this finding and described a higher abundance of epilithic biofilms in the lower intertidal zone than the upper zone due to desiccation and UV light (Aleem 1950, Castenholz 1963, Underwood 1984, Thompson et al. 2004). Accordingly, the gradients of abiotic factors in the intertidal may promote adaptation to these stringent conditions and, as a consequence, increase the habitat specialists (Logares et al. 2013). Thus, restricted distribution of taxa in the intertidal zone may be more common than generally accepted and differences between micro- and microorganisms may not be substantial. This also suggests an imprint of marine over ‘terrestrial’ origin of intertidal microorganisms, as it occurs in rocky shore macro-organisms.

Co-occurrence networks of macro and microorganisms showed similar levels of connectivity and transitivity between the clusters despite the large differences in absolute richness, indicative of non-random clustering within the networks (Röttjers and Faust 2018); consequently, these networks had modules (cluster of Species/OTUs highly connected) corresponding roughly with tidal levels. Those clusters of species were generally more connected than the networks as a whole. Freilich et al. (2018) observe that co-occurrence networks might represent niche preferences of component species more than they reflect specific biotic interactions. Therefore, networks constructed from known interactions (i.e., consumption) are not directly comparable to co-occurrence networks. Also, Co-occurrence networks of microorganisms are structured by environmental heterogeneity (e.g., pH, aridity, net primary productivity in soil (Delgado-Baquerizo et al. 2018), and depth in a marine system (Cram et al. 2015)). With this caveat in mind, the fact remains that networks of micro-and macro-organisms show several remarkably similar attributes and some important differences across the tidal gradient.

Highly positive associations were observed for species restricted to the high intertidal zone (Freilich et al. 2018), whereas highly positive correlations were observed for OTUs limited to the low intertidal zone. The low intertidal zone is generally expected to be the benign end of the environmental stress gradient for organisms of marine origin, which include both micro and macroorganisms. Still, the resulting co-occurrence patterns of positive associations differ between both networks. This supports that communities of macro-and microorganism respond in different ways to the same environmental forcing due possibly to differences in their resiliency to Ambiental stress, different mediating effects of biotic interactions, and dormancy in microbial communities (Mestre and Höfer 2021). But, despite orders of magnitude differences in richness and size, the macro and micro-communities respond to stress gradients, giving rise to specific zonation patterns in the intertidal rocky shore.

## Supporting information

Supplemental_AppendixS1

Supplemental_AppendixS2

Supplemental_AppendixS3

Supplemental_AppendixS4

## Acknowledgements

We thank Sergio Celis at ATIKA Ltda, André Comeau at CGEB-IMR. We are also indebted to many students and research assistants at ECIM who collaborated with us in the field and laboratory. Funding for these studies and for international collaboration was provided by CONICYT (ANID) – National PhD scholarship Program 2016 to CMAB, and by Fondecyt grants No. 1160289 and 1200636 to SAN, and No. 1171259 to RDI. Complementary funding was provided by ANID PIA/BASAL FB0002 to SAN.

## Notes

### Competing Interest Statement

The authors have declared no competing interest.

