## Supplemental_AppendixS1 for "Microbial communities network structure across strong environmental gradients: How do they compare to macroorganisms?"

**Appendix S1**

**Table S1.** Description of intertidal zone surveys of macroorganisms along sites in central Chile. The majority of surveys took place in 2003 (321 surveys). Additional sampling took place in 2004 (124 surveys), 1999 (43 surveys), and 1998 (85 surveys). The surveys included 102 high intertidal, 267 mid intertidal, and 247 low intertidal quadrats. The study site for microbial zonation was ECIMN, 33º 50' S, 71º 63' W.

| YEAR | SOUTH LAT | WEST LONG | SITE | TOTAL SURVEYS BY YEAR |
|---|---|---|---|---|
| 1998 | 33.50 | 71.63 | ECIMN | 85 |
| 1998 | 33.50 | 71.63 | ECIMN | |
| 1998 | 33.50 | 71.63 | ECIMN | |
| 1998 | 33.51 | 71.63 | LCRUC | |
| 1998 | 33.51 | 71.63 | LCRUC | |
| 1998 | 33.51 | 71.63 | LCRUC | |
| 1999 | 33.50 | 71.63 | ECIMN | 43 |
| 1999 | 33.51 | 71.63 | LCRUC | |
| 2003 | 33.42 | 71.70 | PTRLN | 321 |
| 2003 | 33.43 | 71.71 | PTRLS | |
| 2003 | 33.45 | 71.68 | TABO | |
| 2003 | 33.49 | 71.64 | SAL | |
| 2003 | 33.50 | 71.63 | ECIMN | |
| 2003 | 33.51 | 71.63 | LCRUC | |
| 2003 | 33.55 | 71.61 | CART | |
| 2003 | 33.56 | 71.62 | PELA | |
| 2003 | 33.59 | 71.62 | SAN | |
| 2004 | 33.45 | 71.68 | TABO | 124 |
| 2004 | 33.50 | 71.63 | ECIMN | |
| 2004 | 33.50 | 71.63 | ECIMN | |
| 2004 | 33.51 | 71.63 | LCRUC | |
| 2004 | 33.55 | 71.61 | CART | |
| 2004 | 33.55 | 71.61 | CART | |

### Appendix S1

**Table S2.** Description and main characteristics of the intertidal zones and control samples of the microbial community (16S rRNA sequencing).

| # | ID | Treatment | Sample Extracted ($cm^2$) | ng dDNA/ uL | Number of reads | Number of OTUs | Number of reads after rarefaction | Number of OTUs after rarefaction |
|---|---|---|---|---|---|---|---|---|
| 1 | 560_HighZone | High Zone | 22826 | 63.6 | 56862 | 907 | 16888 | 848 |
| 2 | 561_HighZone | High Zone | 22139 | 27.6 | 49140 | 980 | 16888 | 937 |
| 3 | 568_HighZone | High Zone | 26536 | 30.4 | 38831 | 746 | 16888 | 722 |
| 4 | 569_HighZone | High Zone | 25057 | 48.2 | 59919 | 958 | 16888 | 926 |
| 5 | 570_HighZone | High Zone | 35562 | 24.0 | 53583 | 666 | 16888 | 644 |
| 6 | 577_HighZone | High Zone | 32009 | 41.0 | 72215 | 792 | 16888 | 713 |
| 7 | 578_HighZone | High Zone | 29780 | 47.2 | 60989 | 556 | 16888 | 499 |
| 8 | 579_HighZone | High Zone | 26944 | 33.4 | 88915 | 665 | 16888 | 571 |
| 9 | 554_MiddleZone | Middle Zone | 18428 | 98.6 | 92795 | 1188 | 16888 | 989 |
| 10 | 562_MiddleZone | Middle Zone | 20842 | 41.8 | 80676 | 877 | 16888 | 759 |
| 11 | 563_MiddleZone | Middle Zone | 22313 | 32.2 | 57616 | 749 | 16888 | 694 |
| 12 | 564_MiddleZone | Middle Zone | 24636 | 45.6 | 59340 | 884 | 16888 | 816 |
| 13 | 571_MiddleZone | Middle Zone | 15625 | 94.8 | 31309 | 893 | 16888 | 872 |
| 14 | 572_MiddleZone | Middle Zone | 21277 | 110.0 | 29862 | 881 | 16888 | 864 |
| 15 | 580_MiddleZone | Middle Zone | 25540 | 97.2 | 34865 | 895 | 16888 | 872 |
| 16 | 582_MiddleZone | Middle Zone | 24194 | 30.8 | 71677 | 1316 | 16888 | 1190 |
| 17 | 556_LowZone | Low Zone | 19953 | 52.8 | 191992 | 2275 | 16888 | 1775 |
| 18 | 557_LowZone | Low Zone | 21375 | 23.8 | 71386 | 1032 | 16888 | 989 |
| 19 | 566_LowZone | Low Zone | 17688 | 19.9 | 16888 | 392 | 16888 | 392 |
| 20 | 574_LowZone | Low Zone | 17302 | 35.2 | 62217 | 1481 | 16888 | 1359 |
| 21 | 583_LowZone | Low Zone | 18560 | 68.0 | 52117 | 1367 | 16888 | 1299 |
| 22 | 584_LowZone | Low Zone | 22465 | 13.5 | 60638 | 1108 | 16888 | 1040 |
| 23 | 585_LowZone | Low Zone | 26207 | 26.8 | 45649 | 1275 | 16888 | 1198 |
| 24 | IROCA | Control Rock Surface | 21657 | 3.7 | 41564 | 304 | 16888 | 299 |
| 25 | IIROCA | Control Rock Surface | 22202 | 3.4 | 42828 | 330 | 16888 | 326 |
| 26 | IIIROCA | Control Rock Surface | 18988 | 2.1 | 32654 | 289 | 16888 | 287 |
| 27 | IVROCA | Control Rock Surface | 17624 | 4.5 | 18342 | 261 | 16888 | 261 |
| 28 | VROCA | Control Rock Surface | 21407 | 2.0 | 22424 | 257 | 16888 | 256 |

### Appendix S1

**Figure S1**. Graphic representation of zonation of the principal Orders of **A)** Macroorganisms and **B)** Microorganisms across the High, Middle and Low intertidal zones. The coefficients of variation (CV%) across tides for each Order of C) macro- and D) micro-organisms are shown below. The line across the bars represents the mean CV%.

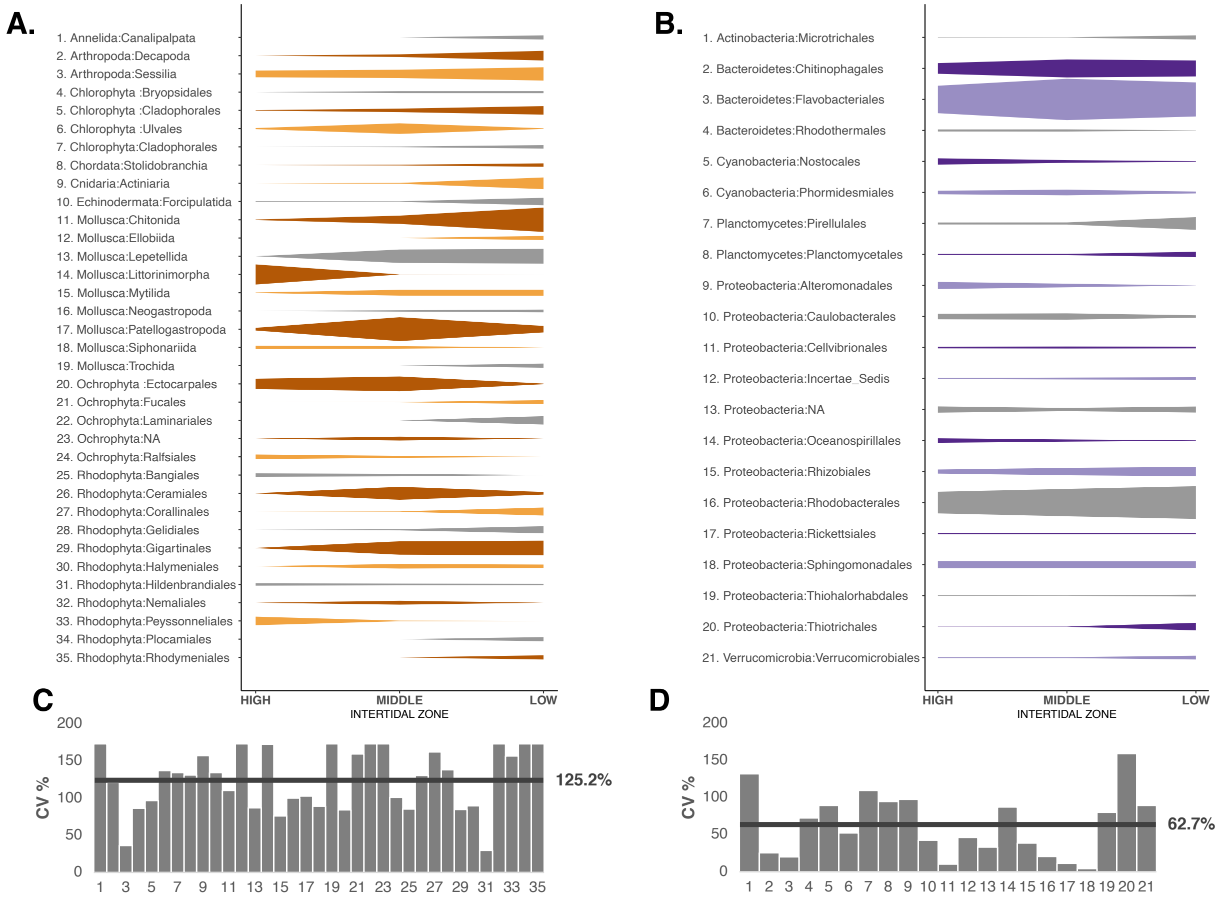

**Appendix S1**

**Table S3.** Cover of sessile species or density of mobile macroalgal and invertebrate species at the a) high, b) middle, and c) low intertidal zone. Data are averages per tidal zone from all surveys described in Table S1.

**A) Mobile species (Density)**

| # | Species | Organism type | Motility | High Average | Mid Average | Low Average | Total All surveys | % All surveys |
|---|---|---|---|---|---|---|---|---|
| 1 | *Acanthopleura echinata* | Chiton | Mobile | 0.00 | 0.06 | 2.74 | 692 | 0.20 |
| 2 | *Chaetopleura peruviana* | Chiton | Mobile | 0.00 | 0.00 | 0.84 | 208 | 0.06 |
| 3 | *Chiton cummingii* | Chiton | Mobile | 0.00 | 0.04 | 0.00 | 12 | <0.001 |
| 4 | *Chiton granosus* | Chiton | Mobile | 2.55 | 14.20 | 4.23 | 5096 | 1.44 |
| 5 | *Chiton latus* | Chiton | Mobile | 0.00 | 0.03 | 0.13 | 40 | 0.01 |
| 6 | *Tonicia chilensis* | Chiton | Mobile | 0.00 | 0.00 | 1.13 | 280 | 0.08 |
| 7 | *Tonicia elegans* | Chiton | Mobile | 0.00 | 0.01 | 0.19 | 52 | 0.01 |
| 8 | *Tonicia spp* | Chiton | Mobile | 0.00 | 0.01 | 0.18 | 48 | 0.01 |
| 9 | *Lottia orbigny* | Limpet | Mobile | 0.00 | 0.03 | 0.02 | 12 | <0.001 |
| 10 | *Scurria spp.* | Limpet | Mobile | 0.00 | 0.16 | 0.00 | 44 | 0.01 |
| 11 | *Fissurella costata* | Limpet | Mobile | 0.00 | 0.00 | 0.32 | 80 | 0.02 |
| 12 | *Fissurella crassa* | Limpet | Mobile | 0.00 | 2.55 | 0.57 | 820 | 0.23 |
| 13 | *Fissurella cummingii* | Limpet | Mobile | 0.00 | 0.13 | 0.13 | 68 | 0.02 |
| 14 | *Fissurella limbata* | Limpet | Mobile | 0.00 | 0.67 | 1.91 | 652 | 0.18 |
| 15 | *Fissurella maxima* | Limpet | Mobile | 0.00 | 0.00 | 0.10 | 24 | 0.01 |
| 16 | *Fissurella picta* | Limpet | Mobile | 0.00 | 0.23 | 0.05 | 74 | 0.02 |
| 17 | *Fissurella pulchra* | Limpet | Mobile | 0.00 | 0.25 | 0.00 | 68 | 0.02 |
| 18 | *Scurria araucana* | Limpet | Mobile | 3.45 | 18.87 | 1.47 | 5753 | 1.62 |
| 19 | *Scurria ceciliana* | Limpet | Mobile | 3.96 | 15.30 | 0.65 | 4649 | 1.31 |
| 20 | *Scurria variabilis* | Limpet | Mobile | 19.45 | 287.58 | 1.25 | 79076 | 22.30 |
| 21 | *Scurria plana* | Limpet | Mobile | 0.00 | 1.73 | 0.02 | 466 | 0.13 |
| 22 | *Scurria scurra* | Limpet | Mobile | 0.00 | 0.52 | 4.32 | 1208 | 0.34 |
| 23 | *Scurria zebrina* | Limpet | Mobile | 0.71 | 2.04 | 0.71 | 793 | 0.22 |
| 24 | *Tegula atra* | Snail | Mobile | 0.00 | 0.00 | 8.39 | 2072 | 0.58 |
| 25 | *Austrolittorina araucana* | Snail | Mobile | 2269.29 | 0.87 | 0.02 | 231704 | 65.33 |
| 26 | *Echinolittorina peruviana* | Snail | Mobile | 66.47 | 0.34 | 0.00 | 6872 | 1.94 |
| 27 | *Trimusculus peruvianus* | pulmonate Limpet | Mobile | 0.00 | 0.00 | 0.03 | 8 | <0.001 |
| 28 | *Concholepas concholepas* | Snail | Mobile | 0.00 | 0.57 | 1.15 | 436 | 0.12 |
| 29 | *Siphonaria lessoni* | pulmonate snail | Mobile | 40.82 | 29.76 | 0.11 | 12139 | 3.42 |
| 30 | *Taliepus dentatus* | Crab | Mobile | 0.00 | 0.00 | 0.05 | 12 | <0.001 |
| 31 | *Acanthocyclus gayi* | Crab | Mobile | 0.00 | 0.45 | 0.60 | 268 | 0.08 |
| 32 | *Acanthocyclus hassleri* | Crab | Mobile | 0.00 | 0.09 | 0.31 | 100 | 0.03 |
| 33 | *Heliaster helianthus* | Sea star | Mobile | 0.16 | 0.10 | 0.70 | 216 | 0.06 |
| 34 | *Stichaster striatus* | Sea star | Mobile | 0.00 | 0.03 | 2.35 | 588 | 0.17 |
| 35 | *Phragmatopoma sp* | polychaete worm | Mobile | 0.00 | 0.00 | 0.06 | 15 | <0.001 |

### B) Sessile species (Cover)

| # | Species | Organism type | Motility | High Average | Mid Average | Low Average | Total All surveys | % All surveys |
|---|---|---|---|---|---|---|---|---|
| 1 | *Brachidontes granulata* | Mussel | Sessile | 0.00 | 0.07 | 0.18 | 63 | 0.10 |
| 2 | *Perumytilus purpuratus* | Mussel | Sessile | 3.94 | 27.91 | 1.40 | 8199 | 12.39 |
| 3 | *Semimytilus algosus* | Mussel | Sessile | 0.00 | 0.03 | 0.09 | 30 | 0.05 |
| 4 | *Jhelius cirratus* | Barnacle | Sessile | 40.24 | 8.71 | 0.01 | 6432 | 9.72 |
| 5 | *Nothochthamalus scabrosus* | Barnacle | Sessile | 0.34 | 5.08 | 0.03 | 1399 | 2.11 |
| 6 | *Austromegabalanus psittacus* | Barnacle | Sessile | 0.00 | 0.00 | 0.02 | 4 | 0.01 |
| 7 | *Balanus spp* | Barnacle | Sessile | 0.00 | 0.00 | 0.10 | 24 | 0.04 |
| 8 | *Nothobalanus flosculus* | Barnacle | Sessile | 0.01 | 0.55 | 8.14 | 2159 | 3.26 |
| 9 | *Balanus laevis* | Barnacle | Sessile | 0.00 | 0.13 | 0.05 | 48 | 0.07 |
| 10 | *Anthotoe spp.* | Anemone | Sessile | 0.00 | 0.00 | 0.02 | 6 | 0.01 |
| 11 | *Isolauctis spp.* | Anemone | Sessile | 0.00 | 0.00 | 0.00 | 1 | <0.001 |
| 12 | *Phymactis spp.* | Anemone | Sessile | 0.00 | 0.31 | 1.30 | 403 | 0.61 |
| 13 | *Pyura chilensis* | sea squirt | Sessile | 0.00 | 0.01 | 0.04 | 11 | 0.02 |
| 14 | *Centroceras spp.* | Macroalgae | Sessile | 0.00 | 0.08 | 0.07 | 38 | 0.06 |
| 15 | *Ceramium spp.* | Macroalgae | Sessile | 0.02 | 0.83 | 0.10 | 248 | 0.37 |
| 16 | *Chaetomorpha_spp* | Macroalgae | Sessile | 0.00 | 0.01 | 0.00 | 4 | 0.01 |
| 17 | *Cladophora spp.* | Macroalgae | Sessile | 0.01 | 0.01 | 0.07 | 20 | 0.03 |
| 18 | *Ectocarpus silicosus* | Macroalgae | Sessile | 0.01 | 0.00 | 0.00 | 1 | <0.001 |
| 19 | *Polysiphonia_spp* | Macroalgae | Sessile | 0.03 | 3.92 | 0.36 | 1140 | 1.72 |
| 20 | *Rama novae-zelandiae* | Macroalgae | Sessile | 0.00 | 0.03 | 0.45 | 117 | 0.18 |
| 21 | *Rhizoclonium cilindricum* | Macroalgae | Sessile | 0.00 | 0.00 | 0.02 | 6 | 0.01 |
| 22 | *Enteromorpha compressa* | Macroalgae | Sessile | 0.00 | 0.02 | 0.00 | 6 | 0.01 |
| 23 | *Pyropia _spp* | Macroalgae | Sessile | 0.84 | 0.59 | 0.03 | 251 | 0.38 |
| 24 | *Scythosiphon lomentaria* | Macroalgae | Sessile | 0.00 | 0.05 | 0.00 | 14 | 0.02 |
| 25 | *Ulva rigida* | Macroalgae | Sessile | 0.30 | 1.13 | 0.17 | 377 | 0.57 |
| 26 | *Petalonia fascia* | Macroalgae | Sessile | 0.01 | 0.14 | 0.03 | 45 | 0.07 |
| 27 | *Sacorthalia sp* | Macroalgae | Sessile | 0.00 | 0.05 | 0.01 | 17 | 0.03 |
| 28 | *Adenocystis utricularis* | Macroalgae | Sessile | 0.00 | 0.00 | 0.00 | 1 | <0.001 |
| 29 | *Ahnfeltiopsis spp.* | Macroalgae | Sessile | 0.00 | 0.00 | 0.01 | 3 | <0.001 |
| 30 | *Chondrus canaliculatus* | Macroalgae | Sessile | 0.00 | 0.00 | 0.02 | 5 | 0.01 |
| 31 | *Codium dimorphum* | Macroalgae | Sessile | 0.00 | 0.90 | 0.94 | 471 | 0.71 |
| 32 | *Colpomenia sinuosa* | Macroalgae | Sessile | 0.00 | 0.04 | 0.00 | 10 | 0.02 |
| 33 | *Gelidium rex* | Macroalgae | Sessile | 0.00 | 0.00 | 0.40 | 100 | 0.15 |
| 34 | *Gelidium spp.* | Macroalgae | Sessile | 0.02 | 2.86 | 7.21 | 2545 | 3.85 |
| 35 | *Grateloupia* | Macroalgae | Sessile | 0.00 | 0.00 | 0.02 | 5 | 0.01 |
| 36 | *Gymnogongrus furcellatus* | Macroalgae | Sessile | 0.00 | 0.00 | 0.52 | 129 | 0.19 |
| 37 | *Laurencia chilensis* | Macroalgae | Sessile | 0.00 | 0.05 | 0.00 | 14 | 0.02 |
| 38 | *Mazzaella laminarioides* | Macroalgae | Sessile | 0.66 | 10.78 | 0.07 | 2961 | 4.48 |
| 39 | *Nothogenia* | Macroalgae | Sessile | 0.00 | 0.04 | 0.00 | 11 | 0.02 |
| 40 | *Petroglossum spp.* | Macroalgae | Sessile | 0.00 | 0.00 | 0.00 | 1 | <0.001 |
| 41 | *Plocamium cartilageneum* | Macroalgae | Sessile | 0.00 | 0.00 | 0.02 | 4 | 0.01 |
| 42 | *Prionitis spp* | Macroalgae | Sessile | 0.00 | 0.01 | 0.00 | 4 | 0.01 |
| 43 | *Rhodymenia sp* | Macroalgae | Sessile | 0.00 | 0.00 | 0.09 | 23 | 0.03 |
| 44 | *Schottera nicaensis* | Macroalgae | Sessile | 0.00 | 0.01 | 0.08 | 22 | 0.03 |
| 45 | *Durvillaea antarctica* | Macroalgae | Sessile | 0.00 | 0.36 | 6.22 | 1632 | 2.47 |
| 46 | *Lessonia nigrescens* | Macroalgae | Sessile | 0.00 | 0.01 | 84.91 | 20976 | 31.70 |
| 47 | *Macrocystis pyrifera* | Macroalgae | Sessile | 0.00 | 0.00 | 0.05 | 12 | 0.02 |
| 48 | *Corallina offcinalis var. chilensis* | Macroalgae | Sessile | 0.01 | 0.63 | 8.28 | 2216 | 3.35 |
| 49 | *Hildenbrandia lecanelieri* | Macroalgae | Sessile | 6.16 | 4.41 | 3.40 | 2646 | 4.00 |
| 50 | *Lithothamnion spp.* | Macroalgae | Sessile | 0.00 | 0.52 | 41.01 | 10269 | 15.52 |
| 51 | *Ralfsia californica* | Macroalgae | Sessile | 1.02 | 0.44 | 0.02 | 227 | 0.34 |
| 52 | *Ulvella spp.* | Macroalgae | Sessile | 0.09 | 1.20 | 0.05 | 341 | 0.52 |
| 53 | *Peysonella spp.* | Macroalgae | Sessile | 3.78 | 0.26 | 0.00 | 456 | 0.69 |

### Appendix S1

**Figure S2.** Relative abundance of the microbial community (Bacteria and Archaea) at the levels of Phylum and Class in samples from the high to low the intertidal zones from left to right. The composition in the Control for the rock surface is also shown.

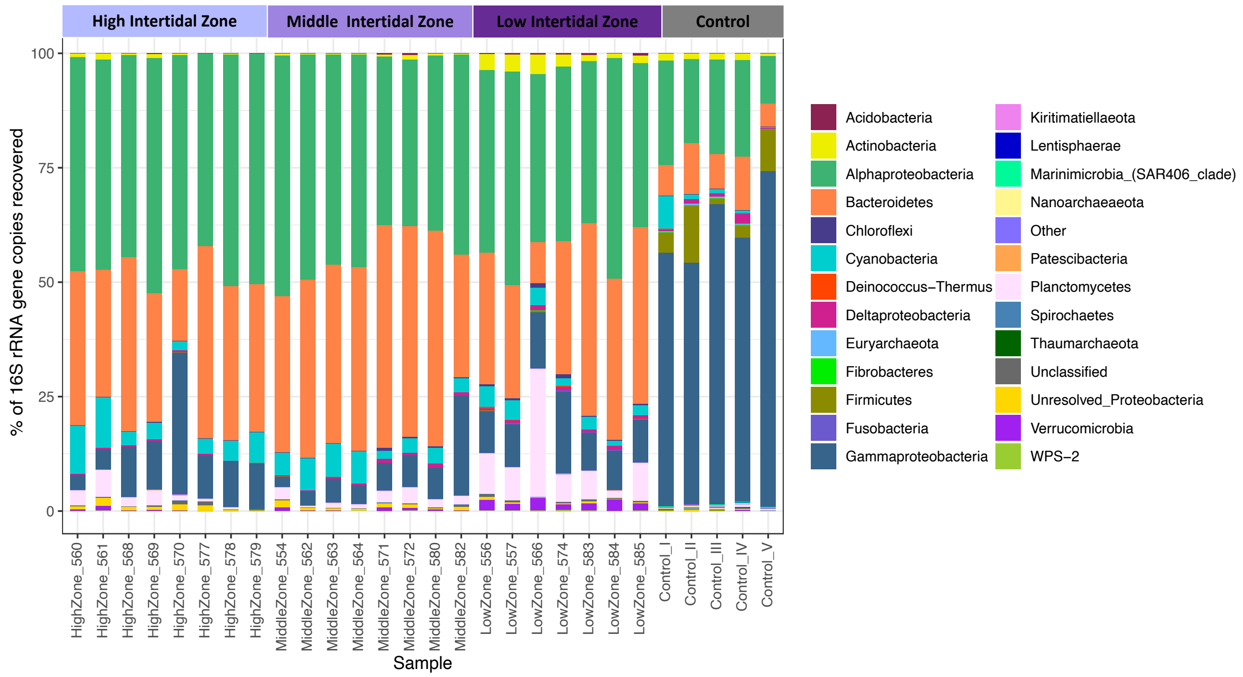


Widely distributed groups in the Chilean rocky intertidal shore belong to Alphaproteobacteria, Gammaproteobacteria, Bacteroidetes, Cyanobacteria, Planctomycetes, Actinobacteria, Verrucomicrobia; these groups were previously described in biofilms of similar systems (Lee et al. 2014, Taylor et al. 2014, Tan et al. 2015, Kerfahi et al. 2020), but this studies also described groups that we do not find abundant, like Chloroflexi, Gemmatinomadetes, Chlorobi (Kerfahi et al. 2020), Deltaproteobacteria and Firmicutes (Tan et al. 2015). All this groups have been widely described on marine biofilms (Dang and Lovell 2016).

### Appendix S1

**Table S3. Main t**axa of microorganisms and their abundance expressed as percentage of total found in each tidal zone and their contribution to the total number of reads (% Total) The representation of these taxa in the control rock surface is also shown. Figures in red correspond to the most abundant taxa of major microbial phyla, i.e., relative abundance > 1% (Pedrós-Alió, 2012).

| TAXA | High Zone % | Middle Zone % | Low Zone % | Rock surface control % |
|---|---|---|---|---|
| Acidobacteria | 0.03 | 0.10 | 0.29 | 0.02 |
| Actinobacteria | 0.55 | 0.46 | 2.56 | 1.27 |
| Alphaproteobacteria | 47.16 | 43.56 | 39.99 | 18.62 |
| Bacteroidetes | 31.44 | 40.09 | 29.65 | 8.49 |
| Chloroflexi | 0.05 | 0.21 | 0.43 | 0.03 |
| Cyanobacteria | 5.61 | 4.76 | 2.95 | 2.02 |
| Deinococcus-Thermus | 0.01 | 0.00 | 0.13 | 0.06 |
| Deltaproteobacteria | 0.25 | 0.50 | 0.70 | 0.83 |
| Euryarchaeota | 0.00 | 0.00 | 0.00 | 0.28 |
| Fibrobacteres | 0.00 | 0.00 | 0.00 | 0.03 |
| Firmicutes | 0.04 | 0.04 | 0.17 | 6.06 |
| Fusobacteria | 0.02 | 0.01 | 0.16 | 0.00 |
| Gammaproteobacteria | 11.21 | 7.01 | 10.63 | 60.86 |
| Kiritimatiellaeota | 0.00 | 0.00 | 0.04 | 0.01 |
| Lentisphaerae | 0.01 | 0.00 | 0.01 | 0.06 |
| Marinimicrobia_(SAR406_clade) | 0.00 | 0.01 | 0.00 | 0.24 |
| Nanoarchaeaeota | 0.00 | 0.00 | 0.00 | 0.12 |
| Other | 0.01 | 0.01 | 0.04 | 0.14 |
| Patescibacteria | 0.02 | 0.01 | 0.06 | 0.04 |
| Planctomycetes | 2.17 | 1.96 | 9.52 | 0.34 |
| Spirochaetes | 0.00 | 0.00 | 0.01 | 0.04 |
| Thaumarchaeota | 0.01 | 0.00 | 0.01 | 0.14 |
| Unclassified | 0.34 | 0.20 | 0.34 | 0.10 |
| Unresolved_Proteobacteria | 0.79 | 0.60 | 0.23 | 0.10 |
| Verrucomicrobia | 0.25 | 0.44 | 1.97 | 0.09 |
| WPS-2 | 0.03 | 0.03 | 0.12 | 0.00 |
| **TOTAL** | **100** | **100** | **100** | **100** |
