## Supplemental_AppendixS2 for "Microbial communities network structure across strong environmental gradients: How do they compare to macroorganisms?"

**Appendix S2**

**Figure S1.** Linear regression between the coefficient of variation across tidal zones and the Species/OTUs richness within Orders.

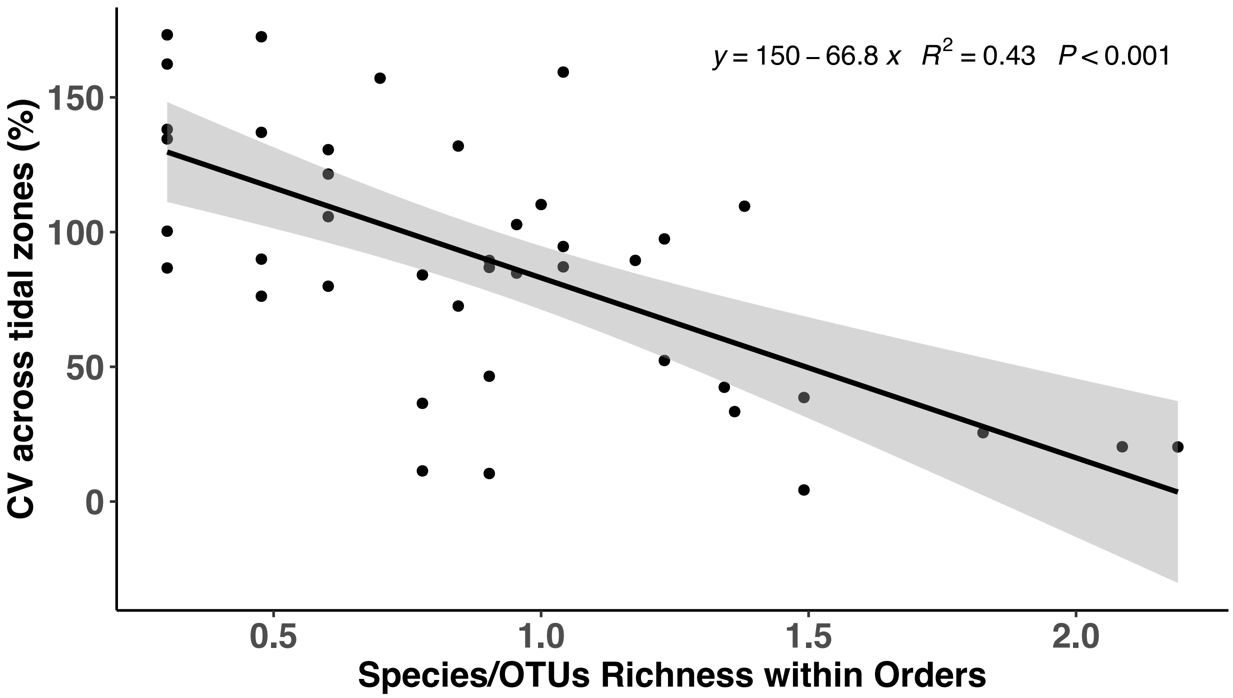

**Appendix S2**

**Figure S2.** Mean **A**) richness and **B**) Shannon index diversity of the microbial communities (N= 6961 OTUs) of the high zone, middle zone, low zone and control of the rock surface (mean + SE). Different letters above bars indicate significant differences (among treatments from **A)** Dunn and **B)** Games-howell test at the experiment wise error rate = 0.

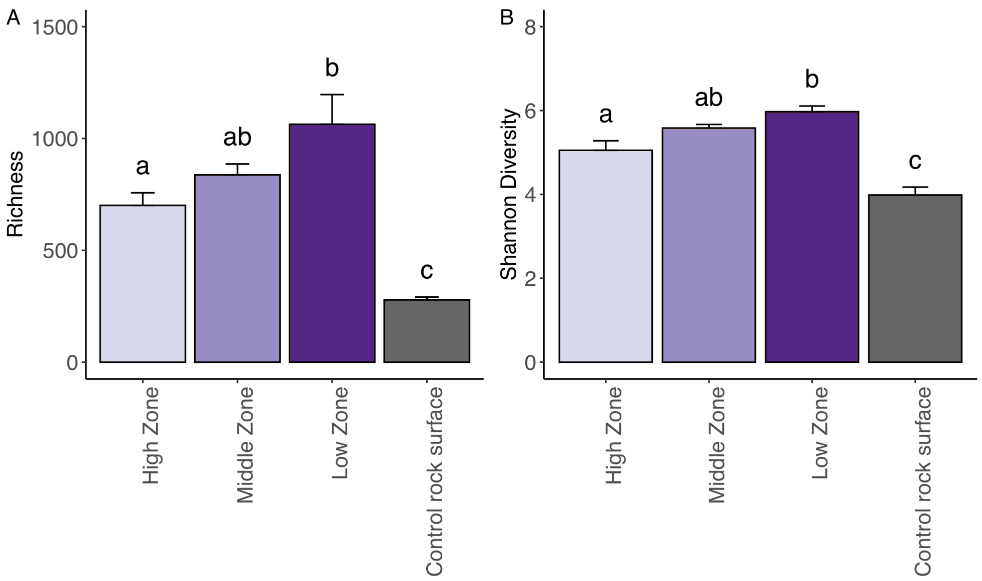
