## Supplemental_AppendixS3 for "Microbial communities network structure across strong environmental gradients: How do they compare to macroorganisms?"

**Appendix S3**

**Figure S1.** Microbial compositional similarity of the different intertidal rocky shore zones. Non-metric multidimensional scaling (NMDS) ordination plots based on Bray-Curtis distances. N= 6961 OTUs. Stress = 0.053. The shapes denote the intertidal zone surrounded by an ellipse showing the 95% confidence interval: (▲) High intertidal zone, (●) Middle intertidal zone, (♦) Low intertidal zone and (■) Control of rock surface.

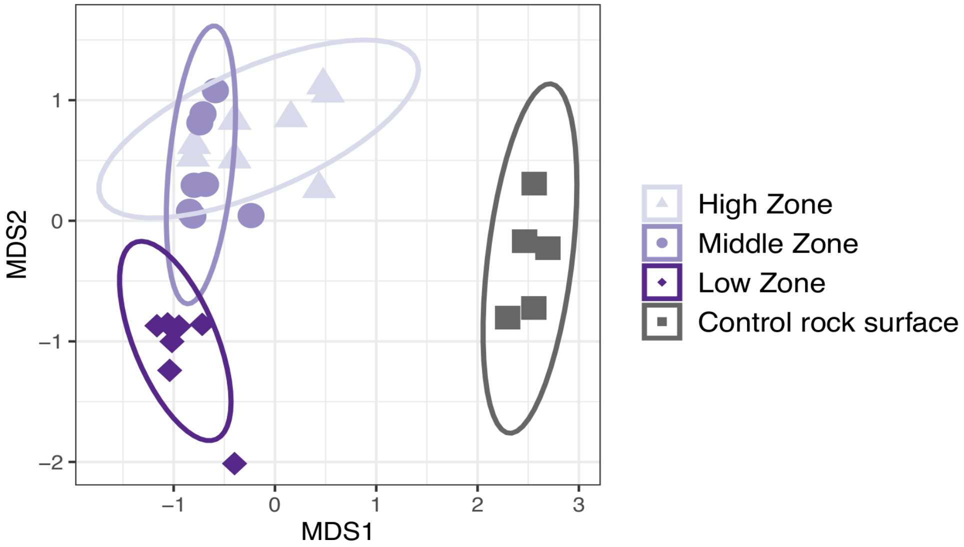


We found statistical differences between all the treatments (PERMANOVA, p=0.0009), and microbial communities of the three intertidal zones were significant different from the microorganisms found in the rock used for biofilm colonization (FDR-corrected pairwise comparisons, High zone: p=0.006 , Middle zone: p=0.012, Low zone: p=0.006).

**Appendix S3**

**Table S1.** False Discovery Rate (FDR) corrected (Benjamini and Hochberg 1995) pairwise comparisons of the macro and microorganisms communities at the high, middle and low intertidal zone. The distances used were (**A,C)** Jaccard and (**B,D)** Bray-Curtis.

### MACROORGANISMS

#### A. Jaccard Distance

| | pairs | Df | Sums Of Sqs | F Model | R2 | p value | p adjusted sig |
|---|---|---|---|---|---|---|---|
| 1 | HighZone vs MiddleZone | 1 | 2.289 | 5.528 | 0.015 | 0.001 | 0.003 * |
| 2 | HighZone vs LowZone | 1 | 1.777 | 4.326 | 0.012 | 0.001 | 0.003 * |
| 3 | MiddleZone vs LowZone | 1 | 0.863 | 2.061 | 0.004 | 0.012 | 0.036 . |

#### B. Bray-Curtis Distance

| | pairs | Df | Sums Of Sqs | F Model | R2 | p value | p adjusted sig |
|---|---|---|---|---|---|---|---|
| 1 | HighZone vs MiddleZone | 1 | 2.289 | 5.528 | 0.015 | 0.001 | 0.003 * |
| 2 | HighZone vs LowZone | 1 | 1.777 | 4.326 | 0.012 | 0.001 | 0.003 * |
| 3 | MiddleZone vs LowZone | 1 | 0.863 | 2.061 | 0.004 | 0.009 | 0.027 . |

### MICROORGANISMS

#### C. Jaccard Distance

| | pairs | Df | Sums Of Sqs | F Model | R2 | p value | p adjusted sig |
|---|---|---|---|---|---|---|---|
| 1 | MiddleZone vs LowZone | 1 | 0.475 | 2.029 | 0.135 | 0.035 | 0.0350 . |
| 2 | MiddleZone vs HighZone | 1 | 1.533 | 6.678 | 0.323 | 0.002 | 0.0045 * |
| 3 | LowZone vs HighZone | 1 | 1.203 | 5.955 | 0.314 | 0.003 | 0.0045 |

#### D. Bray-Curtis Distance

| | pairs | Df | Sums Of Sqs | F Model | R2 | p value | p adjusted sig |
|---|---|---|---|---|---|---|---|
| 1 | MiddleZone vs LowZone | 1 | 0.475 | 2.029 | 0.135 | 0.025 | 0.0250 . |
| 2 | MiddleZone vs HighZone | 1 | 1.532 | 6.678 | 0.323 | 0.001 | 0.0030 * |
| 3 | LowZone vs HighZone | 1 | 1.203 | 5.955 | 0.314 | 0.003 | 0.0045 * |

**Appendix S3**

**Figure S2.** Compositional similarity of the different intertidal rocky shore zones for macroorganisms (N= 88 species) and microorganisms (N= 6,252 OTUs). Metric multidimensional scaling (Principal Coordinates Analysis, PCoA) ordination plots based on (**A**, **C**) Jaccard distances and (**B**, **D**) Bray-Curtis distances. The shapes denote the intertidal zone surrounded by an ellipse showing the 95% confidence interval. Each observation in the graphs is a survey unit (50 x 50 cm quadrats for macroscopic organisms, 3 x 8 cm rock surface coupon for microbes).

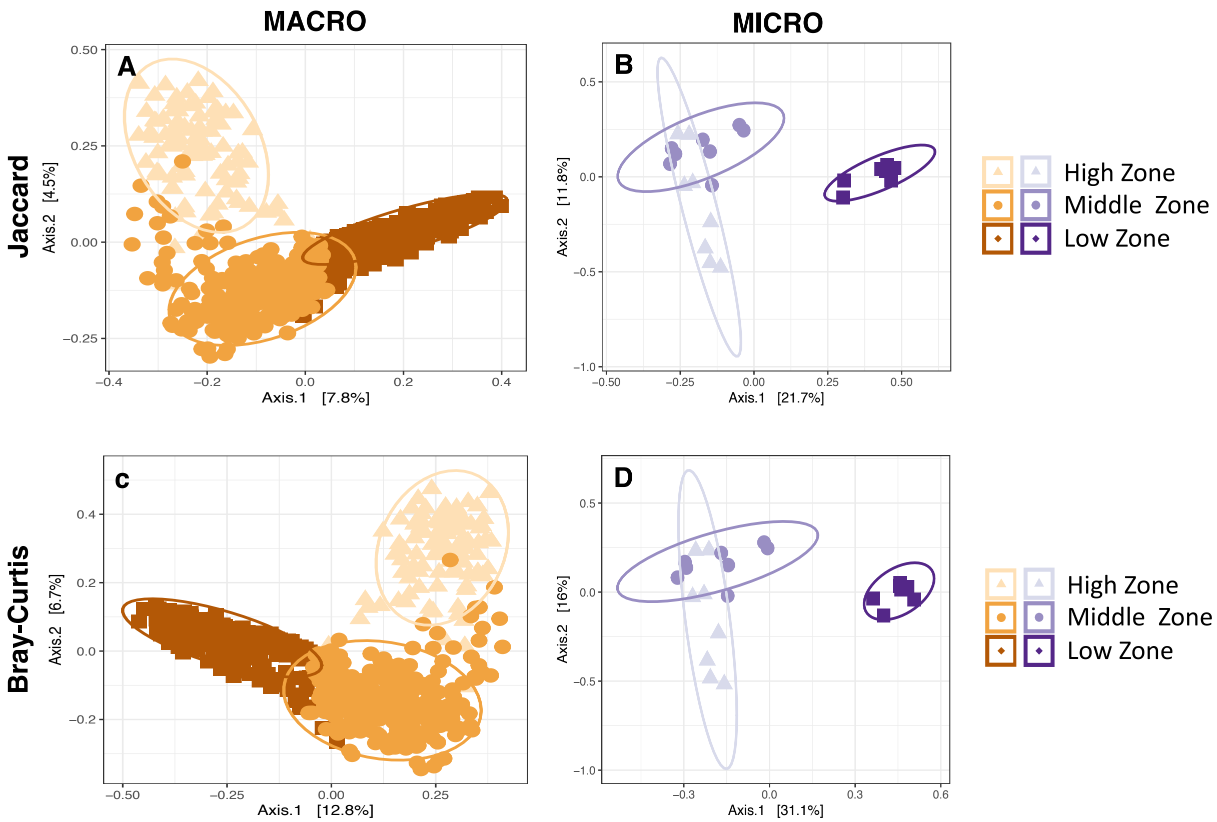

### Appendix S3

**Figure S3.** Mahalanobis Distances for (●) Macro and (●) Micro communities.

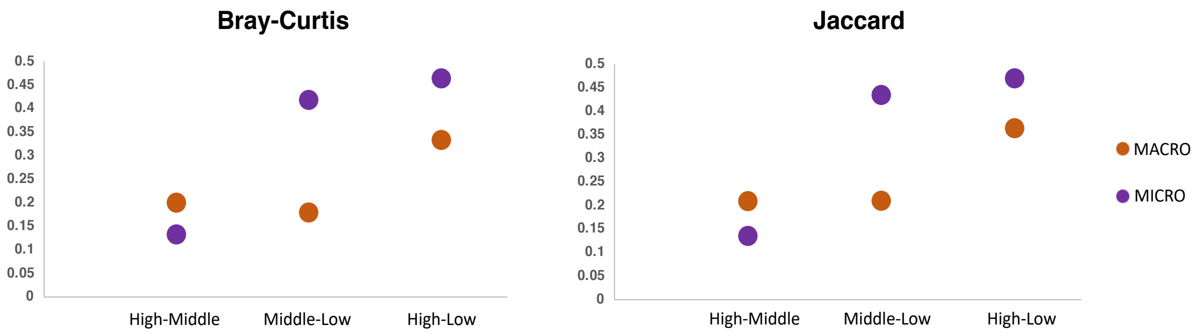
