## Supplemental_AppendixS4 for "Microbial communities network structure across strong environmental gradients: How do they compare to macroorganisms?"

**Appendix S4**

**Table S1.** Strict habitat specialists to a specific intertidal zone, the p-value is shown. **A)** macroorganisms species and **B)** microorganisms OTUs. Only significant value are presented.

### A. Macroorganisms

| Species | pvalue | Intertidal Zone | Phyllum | Class | Order | Family |
|---|---|---|---|---|---|---|
| *acanthopleura echinata* | 0.001 | Low | Mollusca | Polyplacophora | Chitonida | Chitonidae |
| *chaetopleura peruviana* | 0.002 | Low | Mollusca | Polyplacophora | Chitonida | Chaetopleuridae |
| *chiton granosus* | 0.001 | Middle | Mollusca | Polyplacophora | Chitonida | Chitonidae |
| *chiton latus* | 0.024 | Low | Mollusca | Polyplacophora | Chitonida | Chitonidae |
| *tonicia chilensis* | 0.001 | Low | Mollusca | Polyplacophora | Chitonida | Chitonidae |
| *tonicia elegans* | 0.006 | Low | Mollusca | Polyplacophora | Chitonida | Chitonidae |
| *fissurella costata* | 0.002 | Low | Mollusca | Gastropoda | Lepetellida | Fissurellidae |
| *fissurella crassa* | 0.001 | Middle | Mollusca | Gastropoda | Lepetellida | Fissurellidae |
| *fissurella limbata* | 0.001 | Low | Mollusca | Gastropoda | Lepetellida | Fissurellidae |
| *fissurella maxima* | 0.049 | Low | Mollusca | Gastropoda | Lepetellida | Fissurellidae |
| *fissurella puhlcra* | 0.048 | Middle | Mollusca | Gastropoda | Lepetellida | Fissurellidae |
| *scurria araucana* | 0.001 | Middle | Mollusca | Gastropoda | Patellogastropoda | Lottiidae |
| *scurria ceciliana* | 0.001 | Middle | Mollusca | Gastropoda | Patellogastropoda | Lottiidae |
| *scurria variabilis* | 0.001 | Middle | Mollusca | Gastropoda | Patellogastropoda | Lottiidae |
| *scurria plana* | 0.017 | Middle | Mollusca | Gastropoda | Patellogastropoda | Lottiidae |
| *scurria scurra* | 0.001 | Low | Mollusca | Gastropoda | Patellogastropoda | Lottiidae |
| *scurria zebrina* | 0.005 | Middle | Mollusca | Gastropoda | Patellogastropoda | Lottiidae |
| *tegula atra* | 0.001 | Low | Mollusca | Gastropoda | Trochida | Tegulidae |
| *austrolittorina araucana* | 0.001 | High | Mollusca | Gastropoda | Littorinimorpha | Littorinidae |
| *nodolittorina peruviana* | 0.001 | High | Mollusca | Gastropoda | Littorinimorpha | Littorinidae |
| *concholepas concholepas* | 0.002 | Low | Mollusca | Gastropoda | Neogastropoda | Muricidae |
| *siphonaria lessoni* | 0.001 | High | Mollusca | Gastropoda | Siphonariida | Siphonariidae |
| *brachidontes granulata* | 0.001 | Low | Mollusca | Bivalvia | Mytilida | Mytilidae |
| *perumytilus purpuratus* | 0.001 | Middle | Mollusca | Bivalvia | Mytilida | Mytilidae |
| *semimytilus algosus* | 0.034 | Low | Mollusca | Bivalvia | Mytilida | Mytilidae |
| *jhelius cirratus* | 0.001 | High | Arthropoda | Hexanauplia | Sessilia | Chthamalidae |
| *nothochthamalus scabrosus* | 0.001 | Middle | Arthropoda | Hexanauplia | Sessilia | Chthamalidae |

| | | | | | | |
|---|---|---|---|---|---|---|
| *austromegabalanus psittacus* | 0.031 | Low | Arthropoda | Hexanauplia | Sessilia | Balanidae |
| *balanus spp* | 0.012 | Low | Arthropoda | Hexanauplia | Sessilia | Balanidae |
| *nothobalanus flosculus* | 0.001 | Low | Arthropoda | Hexanauplia | Sessilia | Archaeobalanidae |
| *balanus laevis* | 0.021 | Middle | Arthropoda | Hexanauplia | Sessilia | Balanidae |
| *acanthocyclus gayi* | 0.02 | Low | Arthropoda | Malacostraca | Decapoda | Belliidae |
| *acanthocyclus hassleri* | 0.002 | Low | Arthropoda | Malacostraca | Decapoda | Belliidae |
| *heliaster helianthus* | 0.014 | Low | Echinodermata | Asteroidea | Forcipulatida | Heliasteridae |
| *stichaster striatus* | 0.001 | Low | Echinodermata | Asteroidea | Forcipulatida | Stichasteridae |
| *anthotoe spp.* | 0.012 | Low | Cnidaria | Anthozoa | Actiniaria | Sagartiidae |
| *phymactis spp.* | 0.001 | Low | Cnidaria | Anthozoa | Actiniaria | Actiniidae |
| *pyura chilensis* | 0.024 | Low | Chordata | Ascidiacea | Stolidobranchia | Pyuridae |
| *Phragmatopoma sp* | 0.002 | Low | Annelida | Polychaeta | Canalipalpata | Sabellariidae |
| *ceramium spp.* | 0.001 | Middle | Rhodophyta | Florideophyceae | Ceramiales | Ceramiaceae |
| *cladophora spp.* | 0.033 | Low | Chlorophyta | Ulvophyceae | Cladophorales | Cladophoraceae |
| *polysiphonia_spp* | 0.001 | Middle | Rhodophyta | Florideophyceae | Ceramiales | Rhodomelaceae |
| *enteromorpha compressa* | 0.021 | Middle | Chlorophyta | Ulvophyceae | Ulvales | Ulvaceae |
| *porphyra_spp* | 0.001 | High | Rhodophyta | Bangiophyceae | Bangiales | Bangiaceae |
| *scythosiphon lomentaria* | 0.005 | Middle | Ochrophyta | Phaeophyceae | Ectocarpales | Scytosiphonaceae |
| *ulva rigida* | 0.001 | Middle | Chlorophyta | Ulvophyceae | Ulvales | Ulvaceae |
| *petalonia fascia* | 0.001 | Middle | Ochrophyta | Phaeophyceae | Ectocarpales | Scytosiphonaceae |
| *codium dimorphum* | 0.021 | Low | Chlorophyta | Ulvophyceae | Bryopsidales | Codiaceae |
| *gelidium spp.* | 0.001 | Low | Rhodophyta | Florideophyceae | Gelidiales | Gelidiaceae |
| *gymnogongrus furcellatus* | 0.007 | Low | Rhodophyta | Florideophyceae | Gigartinales | Phyllophoraceae |
| *mazzaella laminarioides* | 0.001 | Middle | Rhodophyta | Florideophyceae | Gigartinales | Gigartinaceae |
| *nothogenia* | 0.002 | Middle | Rhodophyta | Florideophyceae | Nemaliales | Scinaiaceae |
| *plocamium cartilageneum* | 0.039 | Low | Rhodophyta | Florideophyceae | Plocamiales | Plocamiaceae |
| *rhodymenia sp* | 0.002 | Low | Rhodophyta | Florideophyceae | Rhodymeniales | Rhodymeniaceae |
| *schottera nicaensis* | 0.003 | Low | Rhodophyta | Florideophyceae | Gigartinales | Phyllophoraceae |
| *durvillaea antarctica* | 0.001 | Low | Ochrophyta | Phaeophyceae | Fucales | Durvillaeaceae |
| *lessonia nigrescens* | 0.001 | Low | Ochrophyta | Phaeophyceae | Laminariales | Lessoniaceae |
| *corallina offcinalis var. chilensis* | 0.001 | Low | Rhodophyta | Florideophyceae | Corallinales | Corallinaceae |
| *lithothamnion spp.* | 0.001 | Low | Rhodophyta | Florideophyceae | Corallinales | Lithothamniaceae |
| *ralfsia californica* | 0.015 | High | Ochrophyta | Phaeophyceae | Ralfsiales | Ralfsiaceae |
| *ulvella spp.* | 0.001 | Middle | Chlorophyta | Ulvophyceae | Ulvales | Ulvellaceae |
| *peysonella spp.* | 0.001 | High | Rhodophyta | Florideophyceae | Peyssonneliales | Peyssonneliaceae |

### B. Microorganisms

| OTUs | Intertidal Zone | pvalue | Phylum | Class | Order | Family | Genus |
|---|---|---|---|---|---|---|---|
| ASV_8 | Low | 0.029 | Proteobacteria | Alphaproteobacteria | Rhodobacterales | Rhodobacteraceae | Sulfitobacter |
| ASV_9 | High | 0.001 | Proteobacteria | Alphaproteobacteria | Rhodobacterales | Rhodobacteraceae | Sulfitobacter |
| ASV_11 | Middle | 0.003 | Proteobacteria | Alphaproteobacteria | Rhodobacterales | Rhodobacteraceae | Sulfitobacter |
| ASV_18 | High | 0.007 | Bacteroidetes | Bacteroidia | Flavobacteriales | Flavobacteriaceae | Gramella |
| ASV_24 | Low | 0.001 | Proteobacteria | Alphaproteobacteria | Rhodobacterales | Rhodobacteraceae | Litoreibacter |
| ASV_28 | High | 0.005 | Proteobacteria | Alphaproteobacteria | Rhodobacterales | Rhodobacteraceae | Jannaschia |
| ASV_30 | Low | 0.001 | Proteobacteria | Alphaproteobacteria | Rhodobacterales | Rhodobacteraceae | Loktanella |
| ASV_37 | Low | 0.01 | Proteobacteria | Alphaproteobacteria | Rhodobacterales | Rhodobacteraceae | Celeribacter |
| ASV_41 | High | 0.023 | Proteobacteria | Alphaproteobacteria | Rhodobacterales | Rhodobacteraceae | Sulfitobacter |
| ASV_42 | Middle | 0.014 | Bacteroidetes | Bacteroidia | Flavobacteriales | Flavobacteriaceae | Nonlabens |
| ASV_44 | Middle | 0.037 | Proteobacteria | Alphaproteobacteria | Rhodobacterales | Rhodobacteraceae | NA |
| ASV_59 | Middle | 0.019 | Proteobacteria | Alphaproteobacteria | Rhodobacterales | Rhodobacteraceae | NA |
| ASV_61 | High | 0.003 | Proteobacteria | Alphaproteobacteria | Rhodobacterales | Rhodobacteraceae | NA |
| ASV_63 | Middle | 0.003 | Bacteroidetes | Bacteroidia | Flavobacteriales | Flavobacteriaceae | Maribacter |
| ASV_67 | High | 0.016 | Proteobacteria | Alphaproteobacteria | Rhodobacterales | Rhodobacteraceae | NA |
| ASV_71 | Middle | 0.009 | Bacteroidetes | Bacteroidia | Flavobacteriales | Flavobacteriaceae | Dokdonia |
| ASV_73 | High | 0.002 | Proteobacteria | Alphaproteobacteria | Rhodobacterales | Rhodobacteraceae | Roseobacter |
| ASV_74 | High | 0.013 | Proteobacteria | Alphaproteobacteria | Rhodobacterales | Rhodobacteraceae | Loktanella |
| ASV_75 | Low | 0.003 | Bacteroidetes | Bacteroidia | Flavobacteriales | Flavobacteriaceae | Winogradskyella |
| ASV_76 | Low | 0.001 | Bacteroidetes | Bacteroidia | Flavobacteriales | Flavobacteriaceae | Croceitalea |
| ASV_79 | Low | 0.001 | Planctomycetes | Planctomycetacia | Pirellulales | Pirellulaceae | Rhodopirellula |
| ASV_80 | Low | 0.014 | Proteobacteria | Alphaproteobacteria | Rhodobacterales | Rhodobacteraceae | Litoreibacter |
| ASV_81 | Low | 0.005 | Proteobacteria | Alphaproteobacteria | Rhodobacterales | Rhodobacteraceae | Loktanella |
| ASV_94 | Low | 0.003 | Proteobacteria | Alphaproteobacteria | Rhodobacterales | Rhodobacteraceae | Boseongicola |
| ASV_99 | Middle | 0.025 | Proteobacteria | Alphaproteobacteria | Rhodobacterales | Rhodobacteraceae | Loktanella |
| ASV_103 | Low | 0.002 | Proteobacteria | Alphaproteobacteria | Caulobacterales | Hyphomonadaceae | NA |
| ASV_106 | Middle | 0.001 | Bacteroidetes | Bacteroidia | Flavobacteriales | Flavobacteriaceae | NA |
| ASV_109 | Middle | 0.008 | Proteobacteria | Alphaproteobacteria | Caulobacterales | Hyphomonadaceae | Algimonas |
| ASV_110 | Middle | 0.001 | Proteobacteria | Alphaproteobacteria | Rhizobiales | Rhizobiaceae | Pseudahrensia |
| ASV_117 | Low | 0.036 | Actinobacteria | Acidimicrobiia | Microtrichales | Ilumatobacteraceae | Ilumatobacter |
| ASV_124 | Middle | 0.035 | Proteobacteria | Alphaproteobacteria | Rhodobacterales | Rhodobacteraceae | NA |
| ASV_128 | Middle | 0.001 | Bacteroidetes | Bacteroidia | Flavobacteriales | Flavobacteriaceae | Dokdonia |
| ASV_132 | Middle | 0.002 | Proteobacteria | Alphaproteobacteria | Rhodobacterales | Rhodobacteraceae | NA |
| ASV_145 | Low | 0.001 | Proteobacteria | Alphaproteobacteria | Rhodobacterales | Rhodobacteraceae | Planktotalea |
| ASV_148 | Low | 0.001 | Proteobacteria | Gammaproteobacteria | Thiotrichales | Thiotrichaceae | Leucothrix |
| ASV_152 | Middle | 0.032 | Proteobacteria | Alphaproteobacteria | Rhodobacterales | Rhodobacteraceae | Pontivivens |

| | | | | | | | |
|---|---|---|---|---|---|---|---|
| ASV_155 | Low | 0.001 | Actinobacteria | Acidimicrobiia | Microtrichales | Microtrichaceae | NA |
| ASV_156 | Middle | 0.001 | Bacteroidetes | Bacteroidia | Chitinophagales | Saprospiraceae | Lewinella |
| ASV_158 | High | 0.001 | Cyanobacteria | Oxyphotobacteria | Nostocales | Xenococcaceae | Pleurocapsa_PCC-7319 |
| ASV_161 | High | 0.041 | Bacteroidetes | Bacteroidia | Flavobacteriales | Flavobacteriaceae | NA |
| ASV_162 | High | 0.011 | Cyanobacteria | Oxyphotobacteria | Nostocales | Xenococcaceae | Pleurocapsa_PCC-7319 |
| ASV_167 | Low | 0.001 | Bacteroidetes | Bacteroidia | Flavobacteriales | Flavobacteriaceae | Maritimimonas |
| ASV_168 | Low | 0.002 | Planctomycetes | Planctomycetacia | Pirellulales | Pirellulaceae | Rhodopirellula |
| ASV_175 | High | 0.024 | Proteobacteria | Alphaproteobacteria | Rhodobacterales | Rhodobacteraceae | Oceanibulbus |
| ASV_177 | Low | 0.001 | Bacteroidetes | Bacteroidia | Flavobacteriales | Flavobacteriaceae | Maritimimonas |
| ASV_181 | Middle | 0.001 | Proteobacteria | Alphaproteobacteria | Rhizobiales | Rhizobiaceae | Pseudahrensia |
| ASV_182 | High | 0.003 | Proteobacteria | Alphaproteobacteria | Sphingomonadales | Sphingomonadaceae | Erythrobacter |
| ASV_184 | Middle | 0.001 | Proteobacteria | Alphaproteobacteria | Caulobacterales | Hyphomonadaceae | Algimonas |
| ASV_185 | Low | 0.001 | Proteobacteria | Alphaproteobacteria | Rhodobacterales | Rhodobacteraceae | Boseongicola |
| ASV_186 | Low | 0.001 | Proteobacteria | Alphaproteobacteria | Rhodobacterales | Rhodobacteraceae | Marinibacterium |
| ASV_192 | Low | 0.026 | Bacteroidetes | Bacteroidia | Chitinophagales | Saprospiraceae | Lewinella |
| ASV_193 | Middle | 0.037 | Bacteroidetes | Bacteroidia | Flavobacteriales | Flavobacteriaceae | Dokdonia |
| ASV_194 | Low | 0.019 | Proteobacteria | Alphaproteobacteria | Rhodobacterales | Rhodobacteraceae | Boseongicola |
| ASV_197 | Middle | 0.011 | Proteobacteria | Alphaproteobacteria | Rhizobiales | Rhizobiaceae | Pseudahrensia |
| ASV_198 | Middle | 0.019 | Proteobacteria | Gammaproteobacteria | NA | NA | NA |
| ASV_200 | Middle | 0.003 | Proteobacteria | Alphaproteobacteria | Rhizobiales | Rhizobiaceae | Pseudahrensia |
| ASV_205 | Low | 0.003 | Proteobacteria | Alphaproteobacteria | Rhodobacterales | Rhodobacteraceae | NA |
| ASV_207 | High | 0.029 | Proteobacteria | Alphaproteobacteria | Caulobacterales | Parvularculaceae | Parvularcula |
| ASV_209 | Low | 0.001 | Bacteroidetes | Bacteroidia | Flavobacteriales | Flavobacteriaceae | Aquibacter |
| ASV_211 | Low | 0.001 | Proteobacteria | Alphaproteobacteria | Rhizobiales | Rhizobiaceae | Ahrensia |
| ASV_213 | Low | 0.001 | Proteobacteria | Gammaproteobacteria | Thiotrichales | Thiotrichaceae | Leucothrix |
| ASV_219 | Middle | 0.001 | Bacteroidetes | Bacteroidia | Flavobacteriales | Flavobacteriaceae | Polaribacter |
| ASV_221 | Middle | 0.02 | Proteobacteria | Alphaproteobacteria | Rhodobacterales | Rhodobacteraceae | NA |
| ASV_222 | Middle | 0.003 | Proteobacteria | Alphaproteobacteria | Rhodobacterales | Rhodobacteraceae | Octadecabacter |
| ASV_226 | Middle | 0.014 | Bacteroidetes | Bacteroidia | Chitinophagales | Saprospiraceae | NA |
| ASV_228 | Low | 0.001 | Bacteroidetes | Bacteroidia | Flavobacteriales | Flavobacteriaceae | Maribacter |
| ASV_229 | High | 0.025 | Proteobacteria | Alphaproteobacteria | Rhizobiales | Rhizobiaceae | Pseudahrensia |
| ASV_231 | Middle | 0.004 | Bacteroidetes | Bacteroidia | Flavobacteriales | Flavobacteriaceae | Dokdonia |
| ASV_233 | High | 0.014 | Bacteroidetes | Bacteroidia | Flavobacteriales | Flavobacteriaceae | Winogradskyella |
| ASV_247 | Low | 0.003 | Bacteroidetes | Bacteroidia | Flavobacteriales | Flavobacteriaceae | Dokdonia |
| ASV_249 | Low | 0.001 | Proteobacteria | Alphaproteobacteria | Rhodobacterales | Rhodobacteraceae | Loktanella |
| ASV_253 | Middle | 0.002 | Bacteroidetes | Bacteroidia | Flavobacteriales | Flavobacteriaceae | Dokdonia |
| ASV_254 | High | 0.036 | Proteobacteria | Gammaproteobacteria | Pseudomonadales | Moraxellaceae | NA |
| ASV_260 | Middle | 0.004 | Bacteroidetes | Bacteroidia | Flavobacteriales | Flavobacteriaceae | Dokdonia |
| ASV_262 | Middle | 0.014 | Proteobacteria | Alphaproteobacteria | Rhodobacterales | Rhodobacteraceae | NA |

| | | | | | | | |
|---|---|---|---|---|---|---|---|
| ASV_271 | High | 0.01 | Proteobacteria | Gammaproteobacteria | Alteromonadales | Alteromonadaceae | Alteromonas |
| ASV_275 | Low | 0.003 | Bacteroidetes | Bacteroidia | Flavobacteriales | Flavobacteriaceae | Psychroserpens |
| ASV_280 | High | 0.015 | Proteobacteria | Alphaproteobacteria | Rhodobacterales | Rhodobacteraceae | NA |
| ASV_285 | High | 0.017 | Proteobacteria | Gammaproteobacteria | Salinisphaerales | Solimonadaceae | Oceanococcus |
| ASV_288 | Middle | 0.005 | Proteobacteria | Alphaproteobacteria | Rhizobiales | Rhizobiaceae | NA |
| ASV_289 | Low | 0.002 | Proteobacteria | Alphaproteobacteria | Sphingomonadales | Sphingomonadaceae | Sphingorhabdus |
| ASV_290 | High | 0.047 | Proteobacteria | Gammaproteobacteria | Alteromonadales | Alteromonadaceae | Alteromonas |
| ASV_297 | Middle | 0.001 | Bacteroidetes | Bacteroidia | Chitinophagales | Saprospiraceae | Lewinella |
| ASV_300 | Low | 0.001 | Proteobacteria | Alphaproteobacteria | Rhodobacterales | Rhodobacteraceae | Litoreibacter |
| ASV_301 | Middle | 0.007 | Proteobacteria | Alphaproteobacteria | Rhodobacterales | Rhodobacteraceae | Litoreibacter |
| ASV_304 | High | 0.048 | Proteobacteria | Alphaproteobacteria | Rhodobacterales | Rhodobacteraceae | Loktanella |
| ASV_310 | Middle | 0.011 | Cyanobacteria | Oxyphotobacteria | Nostocales | Xenococcaceae | Pleurocapsa_PCC-7319 |
| ASV_313 | High | 0.031 | Bacteroidetes | Bacteroidia | Flavobacteriales | Flavobacteriaceae | Jejudonia |
| ASV_321 | Low | 0.002 | Bacteroidetes | Bacteroidia | Flavobacteriales | Flavobacteriaceae | Lutimonas |
| ASV_328 | Middle | 0.026 | Proteobacteria | Alphaproteobacteria | Caulobacterales | Hyphomonadaceae | Algimonas |
| ASV_330 | High | 0.006 | Proteobacteria | Alphaproteobacteria | Sphingomonadales | Sphingomonadaceae | Altererythrobacter |
| ASV_332 | Middle | 0.001 | Bacteroidetes | Bacteroidia | Flavobacteriales | Flavobacteriaceae | Kordia |
| ASV_335 | Low | 0.001 | Planctomycetes | Planctomycetacia | Pirellulales | Pirellulaceae | Rubripirellula |
| ASV_337 | Low | 0.001 | Bacteroidetes | Bacteroidia | Flavobacteriales | Flavobacteriaceae | NA |
| ASV_339 | Low | 0.001 | Bacteroidetes | Bacteroidia | Flavobacteriales | Flavobacteriaceae | NA |
| ASV_346 | Low | 0.001 | Proteobacteria | Alphaproteobacteria | Rhodobacterales | Rhodobacteraceae | NA |
| ASV_347 | Low | 0.001 | Planctomycetes | Planctomycetacia | Planctomycetales | Rubinisphaeraceae | Fuerstia |
| ASV_349 | High | 0.005 | Proteobacteria | Gammaproteobacteria | Salinisphaerales | Solimonadaceae | Oceanococcus |
| ASV_352 | Low | 0.014 | Bacteroidetes | Bacteroidia | Flavobacteriales | Flavobacteriaceae | Lacinutrix |
| ASV_353 | Middle | 0.003 | Proteobacteria | Alphaproteobacteria | Rhizobiales | Rhizobiaceae | Pseudahrensia |
| ASV_355 | High | 0.009 | Proteobacteria | Alphaproteobacteria | Sphingomonadales | Sphingomonadaceae | Erythrobacter |
| ASV_358 | Middle | 0.001 | Bacteroidetes | Bacteroidia | Flavobacteriales | Flavobacteriaceae | NA |
| ASV_359 | Low | 0.001 | Proteobacteria | Gammaproteobacteria | NA | NA | NA |
| ASV_361 | High | 0.019 | Proteobacteria | Alphaproteobacteria | NA | NA | NA |
| ASV_362 | Low | 0.007 | Bacteroidetes | Bacteroidia | Chitinophagales | Saprospiraceae | Lewinella |
| ASV_365 | Low | 0.001 | Proteobacteria | Alphaproteobacteria | Rhizobiales | Hyphomicrobiaceae | Filomicrobium |
| ASV_367 | Low | 0.001 | Proteobacteria | Alphaproteobacteria | Rhodobacterales | Rhodobacteraceae | Roseobacter |
| ASV_368 | Middle | 0.009 | Bacteroidetes | Bacteroidia | Flavobacteriales | Flavobacteriaceae | Dokdonia |
| ASV_374 | High | 0.008 | Proteobacteria | Gammaproteobacteria | Pseudomonadales | Moraxellaceae | Psychrobacter |
| ASV_376 | High | 0.027 | Proteobacteria | Alphaproteobacteria | Sphingomonadales | Sphingomonadaceae | Sphingorhabdus |
| ASV_377 | Low | 0.001 | Bacteroidetes | Bacteroidia | Flavobacteriales | Flavobacteriaceae | NA |
| ASV_378 | High | 0.009 | Bacteroidetes | Bacteroidia | Flavobacteriales | Flavobacteriaceae | Psychroserpens |
| ASV_380 | High | 0.006 | Proteobacteria | Gammaproteobacteria | NA | NA | NA |
| ASV_384 | Low | 0.002 | Bacteroidetes | Bacteroidia | Flavobacteriales | Flavobacteriaceae | Winogradskyella |

| | | | | | | | |
|---|---|---|---|---|---|---|---|
| ASV_388 | Low | 0.001 | Proteobacteria | Gammaproteobacteria | Thiohalorhabdales | Thiohalorhabdaceae | Granulosicoccus |
| ASV_390 | Low | 0.001 | Planctomycetes | Planctomycetacia | Planctomycetales | NA | NA |
| ASV_392 | Low | 0.005 | Bacteroidetes | Bacteroidia | Flavobacteriales | Flavobacteriaceae | Ulvibacter |
| ASV_393 | Low | 0.001 | Proteobacteria | Alphaproteobacteria | Rhodobacterales | Rhodobacteraceae | NA |
| ASV_394 | Low | 0.001 | Proteobacteria | Alphaproteobacteria | Rhizobiales | Rhizobiales_Incertae_Sedis | Anderseniella |
| ASV_397 | Middle | 0.047 | Bacteroidetes | Bacteroidia | Flavobacteriales | Flavobacteriaceae | NA |
| ASV_401 | Middle | 0.028 | Proteobacteria | Alphaproteobacteria | Rhodobacterales | Rhodobacteraceae | NA |
| ASV_406 | High | 0.022 | Bacteroidetes | Bacteroidia | Chitinophagales | Saprospiraceae | NA |
| ASV_410 | Low | 0.001 | Proteobacteria | Alphaproteobacteria | Rhodobacterales | Rhodobacteraceae | Roseovarius |
| ASV_416 | Low | 0.003 | Planctomycetes | Planctomycetacia | Pirellulales | Pirellulaceae | Rhodopirellula |
| ASV_420 | Low | 0.01 | Bacteroidetes | Bacteroidia | Chitinophagales | Saprospiraceae | Lewinella |
| ASV_423 | Low | 0.001 | Proteobacteria | Alphaproteobacteria | Rhodobacterales | Rhodobacteraceae | Amylibacter |
| ASV_425 | Low | 0.002 | Cyanobacteria | Oxyphotobacteria | Thermosynechococcales | Acaryochloridaceae | Acaryochloris_MBIC11017 |
| ASV_432 | Low | 0.002 | Proteobacteria | Alphaproteobacteria | Rhizobiales | Rhizobiaceae | NA |
| ASV_434 | Low | 0.001 | Proteobacteria | Alphaproteobacteria | Rhodobacterales | Rhodobacteraceae | Loktanella |
| ASV_437 | Low | 0.01 | Bacteroidetes | Bacteroidia | Flavobacteriales | Flavobacteriaceae | Olleya |
| ASV_438 | Middle | 0.036 | Bacteroidetes | Bacteroidia | Chitinophagales | Saprospiraceae | Aureispira |
| ASV_442 | Middle | 0.002 | Bacteroidetes | Bacteroidia | Chitinophagales | Saprospiraceae | Lewinella |
| ASV_445 | Low | 0.034 | Proteobacteria | Alphaproteobacteria | Sphingomonadales | Sphingomonadaceae | Altererythrobacter |
| ASV_451 | Middle | 0.003 | Bacteroidetes | Bacteroidia | Chitinophagales | Saprospiraceae | Lewinella |
| ASV_453 | Low | 0.011 | Proteobacteria | Gammaproteobacteria | Gammaproteobacteria_Incertae_Sedis | Unknown_Family | Marinicella |
| ASV_454 | High | 0.001 | Cyanobacteria | Oxyphotobacteria | Nostocales | Xenococcaceae | Pleurocapsa_PCC-7319 |
| ASV_458 | Middle | 0.011 | Bacteroidetes | Bacteroidia | Chitinophagales | Saprospiraceae | NA |
| ASV_461 | Middle | 0.019 | Bacteroidetes | Bacteroidia | Flavobacteriales | Flavobacteriaceae | NA |
| ASV_462 | Low | 0.002 | Verrucomicrobia | Verrucomicrobiae | Verrucomicrobiales | Rubritaleaceae | Roseibacillus |
| ASV_464 | Low | 0.001 | Proteobacteria | Gammaproteobacteria | Thiotrichales | Thiotrichaceae | Leucothrix |
| ASV_467 | Low | 0.013 | Bacteroidetes | Bacteroidia | Chitinophagales | Saprospiraceae | Lewinella |
| ASV_472 | Middle | 0.001 | Proteobacteria | Alphaproteobacteria | Rhodobacterales | Rhodobacteraceae | Pontivivens |
| ASV_476 | High | 0.033 | Bacteroidetes | Bacteroidia | Chitinophagales | Saprospiraceae | NA |
| ASV_477 | Middle | 0.014 | Proteobacteria | Alphaproteobacteria | Rickettsiales | NA | NA |
| ASV_478 | Low | 0.027 | Bacteroidetes | Bacteroidia | Flavobacteriales | Flavobacteriaceae | Psychroserpens |
| ASV_479 | Low | 0.001 | Proteobacteria | Alphaproteobacteria | Rhizobiales | Rhizobiaceae | NA |
| ASV_481 | Middle | 0.001 | Bacteroidetes | Bacteroidia | Flavobacteriales | Flavobacteriaceae | Dokdonia |
| ASV_482 | Low | 0.001 | Bacteroidetes | Bacteroidia | Chitinophagales | Saprospiraceae | NA |
| ASV_483 | Middle | 0.004 | Proteobacteria | Alphaproteobacteria | Rhodobacterales | Rhodobacteraceae | Roseobacter |
| ASV_494 | Low | 0.001 | Planctomycetes | Planctomycetacia | Pirellulales | Pirellulaceae | Blastopirellula |
| ASV_496 | Low | 0.001 | Proteobacteria | Alphaproteobacteria | Rhodobacterales | Rhodobacteraceae | Planktotalea |
| ASV_505 | Low | 0.001 | Bacteroidetes | Bacteroidia | Flavobacteriales | Flavobacteriaceae | Croceitalea |
| ASV_508 | Low | 0.001 | Bacteroidetes | Bacteroidia | Flavobacteriales | Flavobacteriaceae | Maritimimonas |

| | | | | | | | |
|---|---|---|---|---|---|---|---|
| ASV_510 | Middle | 0.004 | Bacteroidetes | Bacteroidia | Flavobacteriales | Flavobacteriaceae | Polaribacter |
| ASV_512 | Low | 0.001 | Proteobacteria | Alphaproteobacteria | Rhodobacterales | Rhodobacteraceae | NA |
| ASV_513 | Low | 0.014 | Proteobacteria | Alphaproteobacteria | Rhizobiales | Rhizobiaceae | Pseudahrensia |
| ASV_514 | Low | 0.001 | Proteobacteria | Alphaproteobacteria | Sphingomonadales | Sphingomonadaceae | NA |
| ASV_515 | High | 0.001 | Proteobacteria | Alphaproteobacteria | Rhodobacterales | Rhodobacteraceae | Planktotalea |
| ASV_516 | Low | 0.001 | Bacteroidetes | Bacteroidia | Chitinophagales | Saprospiraceae | NA |
| ASV_518 | Low | 0.001 | Proteobacteria | Gammaproteobacteria | Thiotrichales | Thiotrichaceae | Leucothrix |
| ASV_519 | Low | 0.002 | Bacteroidetes | Bacteroidia | Chitinophagales | Saprospiraceae | Lewinella |
| ASV_521 | Middle | 0.015 | Bacteroidetes | Bacteroidia | Chitinophagales | Saprospiraceae | NA |
| ASV_527 | Low | 0.004 | Proteobacteria | Alphaproteobacteria | Rhodobacterales | Rhodobacteraceae | Loktanella |
| ASV_528 | Low | 0.001 | Proteobacteria | Alphaproteobacteria | Rhodobacterales | Rhodobacteraceae | Planktotalea |
| ASV_529 | Low | 0.001 | Proteobacteria | Alphaproteobacteria | Rhodobacterales | Rhodobacteraceae | Roseobacter_clade_NAC 11-7_lineage |
| ASV_536 | Middle | 0.019 | Proteobacteria | Alphaproteobacteria | Rhodobacterales | Rhodobacteraceae | NA |
| ASV_538 | Low | 0.007 | Proteobacteria | Gammaproteobacteria | Cellvibrionales | Halieaceae | Marimicrobium |
| ASV_539 | Low | 0.002 | Actinobacteria | Acidimicrobiia | Microtrichales | Ilumatobacteraceae | Ilumatobacter |
| ASV_541 | Low | 0.001 | Planctomycetes | Planctomycetacia | Planctomycetales | NA | NA |
| ASV_543 | Middle | 0.027 | Bacteroidetes | Bacteroidia | Chitinophagales | Saprospiraceae | Aureispira |
| ASV_544 | Middle | 0.021 | Bacteroidetes | Bacteroidia | Chitinophagales | Saprospiraceae | Lewinella |
| ASV_545 | Low | 0.001 | Proteobacteria | Alphaproteobacteria | Rhodobacterales | Rhodobacteraceae | Octadecabacter |
| ASV_550 | Low | 0.033 | Planctomycetes | Planctomycetacia | Pirellulales | Pirellulaceae | Rubripirellula |
| ASV_551 | Low | 0.001 | Proteobacteria | Alphaproteobacteria | Rhizobiales | Rhizobiaceae | Pseudahrensia |
| ASV_554 | High | 0.005 | Proteobacteria | Alphaproteobacteria | Caulobacterales | Hyphomonadaceae | Litorimonas |
| ASV_557 | Middle | 0.003 | Proteobacteria | Alphaproteobacteria | Rhodobacterales | Rhodobacteraceae | NA |
| ASV_559 | Low | 0.001 | Proteobacteria | Alphaproteobacteria | Parvibaculales | PS1_clade | NA |
| ASV_560 | Middle | 0.007 | Proteobacteria | Alphaproteobacteria | Caulobacterales | Hyphomonadaceae | Algimonas |
| ASV_562 | High | 0.002 | Proteobacteria | Alphaproteobacteria | Rhodobacterales | Rhodobacteraceae | Maribius |
| ASV_564 | Middle | 0.039 | Bacteroidetes | Bacteroidia | Flavobacteriales | Flavobacteriaceae | Polaribacter |
| ASV_565 | High | 0.023 | Proteobacteria | Gammaproteobacteria | Alteromonadales | Marinobacteraceae | Marinobacter |
| ASV_566 | Low | 0.001 | Proteobacteria | Alphaproteobacteria | Rhodobacterales | Rhodobacteraceae | NA |
| ASV_569 | Low | 0.001 | Bacteroidetes | Bacteroidia | Flavobacteriales | Flavobacteriaceae | Arcticiflavibacter |
| ASV_578 | Low | 0.002 | Bacteroidetes | Bacteroidia | Flavobacteriales | Flavobacteriaceae | Ulvibacter |
| ASV_579 | Middle | 0.004 | Proteobacteria | Alphaproteobacteria | Rhodobacterales | Rhodobacteraceae | NA |
| ASV_582 | Low | 0.001 | Proteobacteria | Alphaproteobacteria | Sphingomonadales | Sphingomonadaceae | Sphingorhabdus |
| ASV_583 | Middle | 0.019 | Bacteroidetes | Bacteroidia | Flavobacteriales | Flavobacteriaceae | Polaribacter_4 |
| ASV_585 | High | 0.003 | Proteobacteria | Alphaproteobacteria | Rhizobiales | Rhizobiaceae | Aurantimonas |
| ASV_588 | Low | 0.001 | Planctomycetes | Planctomycetacia | Planctomycetales | Rubinisphaeraceae | Fuerstia |
| ASV_589 | Low | 0.001 | Proteobacteria | Alphaproteobacteria | Caulobacterales | Hyphomonadaceae | Hellea |
| ASV_593 | Low | 0.001 | Bacteroidetes | Bacteroidia | NA | NA | NA |
| ASV_594 | Low | 0.003 | Bacteroidetes | Bacteroidia | Chitinophagales | NA | NA |

| | | | | | | | |
|---|---|---|---|---|---|---|---|
| ASV_596 | Low | 0.001 | Proteobacteria | Alphaproteobacteria | NA | NA | NA |
| ASV_597 | High | 0.044 | Bacteroidetes | Bacteroidia | Flavobacteriales | Flavobacteriaceae | Nonlabens |
| ASV_598 | Middle | 0.047 | Proteobacteria | Alphaproteobacteria | Rhizobiales | Rhizobiaceae | Pseudahrensia |
| ASV_602 | Low | 0.002 | NA | NA | NA | NA | NA |
| ASV_608 | Low | 0.001 | Proteobacteria | Alphaproteobacteria | Rhizobiales | Rhizobiaceae | Pseudahrensia |
| ASV_610 | Middle | 0.024 | Bacteroidetes | Bacteroidia | Chitinophagales | Saprospiraceae | Lewinella |
| ASV_611 | High | 0.031 | Proteobacteria | Gammaproteobacteria | Thiohalorhabdales | Thiohalorhabdaceae | Granulosicoccus |
| ASV_612 | Low | 0.001 | Verrucomicrobia | Verrucomicrobiae | Verrucomicrobiales | Rubritaleaceae | Rubritalea |
| ASV_615 | Low | 0.001 | Proteobacteria | Alphaproteobacteria | Rhodobacterales | Rhodobacteraceae | Lentibacter |
| ASV_620 | Middle | 0.029 | Proteobacteria | Alphaproteobacteria | Sphingomonadales | Sphingomonadaceae | Sphingorhabdus |
| ASV_621 | Middle | 0.048 | Proteobacteria | Alphaproteobacteria | Sphingomonadales | Sphingomonadaceae | Altererythrobacter |
| ASV_622 | Low | 0.001 | Bacteroidetes | Bacteroidia | Flavobacteriales | Flavobacteriaceae | Spongiimicrobium |
| ASV_626 | Middle | 0.025 | Cyanobacteria | Oxyphotobacteria | Phormidesmiales | Phormidesmiaceae | Phormidesmis_ANT.LA CV5.1 |
| ASV_628 | High | 0.037 | Cyanobacteria | Oxyphotobacteria | Nostocales | Xenococcaceae | Pleurocapsa_PCC-7319 |
| ASV_629 | Middle | 0.003 | Proteobacteria | Alphaproteobacteria | Caulobacterales | Hyphomonadaceae | Litorimonas |
| ASV_630 | Low | 0.002 | Bacteroidetes | Bacteroidia | Chitinophagales | Saprospiraceae | NA |
| ASV_631 | Low | 0.001 | Proteobacteria | Alphaproteobacteria | Rhizobiales | Rhizobiaceae | Pseudahrensia |
| ASV_635 | Low | 0.001 | Proteobacteria | Alphaproteobacteria | Rhizobiales | Rhizobiaceae | Pseudahrensia |
| ASV_637 | Middle | 0.009 | Proteobacteria | Alphaproteobacteria | Sphingomonadales | Sphingomonadaceae | Altererythrobacter |
| ASV_642 | Low | 0.001 | Bacteroidetes | Bacteroidia | Flavobacteriales | Flavobacteriaceae | NA |
| ASV_645 | Low | 0.001 | Planctomycetes | Planctomycetacia | Pirellulales | Pirellulaceae | Blastopirellula |
| ASV_647 | Middle | 0.012 | Bacteroidetes | Bacteroidia | Flavobacteriales | Flavobacteriaceae | Winogradskyella |
| ASV_648 | Low | 0.001 | Proteobacteria | Alphaproteobacteria | Rhodobacterales | Rhodobacteraceae | Octadecabacter |
| ASV_649 | High | 0.048 | Bacteroidetes | Bacteroidia | Flavobacteriales | Flavobacteriaceae | Pricia |
| ASV_650 | Middle | 0.002 | Proteobacteria | Alphaproteobacteria | Rhodobacterales | Rhodobacteraceae | NA |
| ASV_658 | High | 0.033 | Proteobacteria | Alphaproteobacteria | Rhodobacterales | Rhodobacteraceae | Maribius |
| ASV_660 | Low | 0.001 | Proteobacteria | Gammaproteobacteria | Thiotrichales | Thiotrichaceae | Thiothrix |
| ASV_661 | High | 0.041 | Proteobacteria | Gammaproteobacteria | NA | NA | NA |
| ASV_667 | High | 0.016 | Cyanobacteria | Oxyphotobacteria | Nostocales | Xenococcaceae | Pleurocapsa_PCC-7319 |
| ASV_673 | Low | 0.001 | Proteobacteria | Alphaproteobacteria | Rhodobacterales | Rhodobacteraceae | Loktanella |
| ASV_674 | Low | 0.001 | Proteobacteria | Gammaproteobacteria | Gammaproteobacteria_ Incertae_Sedis | Unknown_Family | Marinicella |
| ASV_675 | Low | 0.002 | Actinobacteria | Acidimicrobiia | Microtrichales | Ilumatobacteraceae | Ilumatobacter |
| ASV_677 | High | 0.006 | Bacteroidetes | Bacteroidia | Flavobacteriales | Flavobacteriaceae | Croceibacter |
| ASV_678 | Middle | 0.043 | Bacteroidetes | Bacteroidia | Flavobacteriales | Flavobacteriaceae | Polaribacter_4 |
| ASV_679 | Low | 0.001 | Planctomycetes | Planctomycetacia | Pirellulales | Pirellulaceae | Blastopirellula |
| ASV_684 | Middle | 0.03 | Bacteroidetes | Rhodothermia | Rhodothermales | Rhodothermaceae | Rubrivirga |
| ASV_687 | Middle | 0.001 | Proteobacteria | Alphaproteobacteria | Caulobacterales | Hyphomonadaceae | Algimonas |
| ASV_690 | Low | 0.001 | Cyanobacteria | Oxyphotobacteria | Phormidesmiales | Phormidesmiaceae | Phormidesmis_ANT.LA CV5.1 |
| ASV_694 | High | 0.005 | Proteobacteria | Alphaproteobacteria | Sphingomonadales | Sphingomonadaceae | Altererythrobacter |

| | | | | | | | |
|---|---|---|---|---|---|---|---|
| ASV_706 | High | 0.006 | Bacteroidetes | Bacteroidia | Flavobacteriales | Flavobacteriaceae | Gramella |
| ASV_718 | Low | 0.001 | Bacteroidetes | Bacteroidia | Flavobacteriales | Flavobacteriaceae | Winogradskyella |
| ASV_724 | High | 0.007 | Bacteroidetes | Bacteroidia | Chitinophagales | Saprospiraceae | NA |
| ASV_725 | Middle | 0.006 | Bacteroidetes | Bacteroidia | Flavobacteriales | Flavobacteriaceae | Dokdonia |
| ASV_726 | Low | 0.001 | Bacteroidetes | Bacteroidia | Chitinophagales | Saprospiraceae | Rubidimonas |
| ASV_738 | Low | 0.001 | Actinobacteria | Acidimicrobiia | Microtrichales | Microtrichaceae | Sva0996_marine_group |
| ASV_740 | Low | 0.001 | Verrucomicrobia | Verrucomicrobiae | Verrucomicrobiales | Rubritaleaceae | Roseibacillus |
| ASV_748 | Low | 0.001 | Proteobacteria | Alphaproteobacteria | Rhizobiales | Rhizobiaceae | NA |
| ASV_751 | Low | 0.001 | Proteobacteria | Gammaproteobacteria | Cellvibrionales | Spongiibacteraceae | NA |
| ASV_756 | Middle | 0.036 | Bacteroidetes | Bacteroidia | Chitinophagales | Saprospiraceae | Lewinella |
| ASV_758 | Low | 0.001 | Proteobacteria | Gammaproteobacteria | Gammaproteobacteria_ Incertae_Sedis | Unknown_Family | Marinicella |
| ASV_759 | Middle | 0.003 | Proteobacteria | Alphaproteobacteria | Sphingomonadales | Sphingomonadaceae | Altererythrobacter |
| ASV_760 | Low | 0.001 | Proteobacteria | Alphaproteobacteria | Rhodobacterales | Rhodobacteraceae | NA |
| ASV_761 | High | 0.004 | Cyanobacteria | Oxyphotobacteria | Nostocales | Xenococcaceae | Pleurocapsa_PCC-7319 |
| ASV_763 | Middle | 0.005 | Bacteroidetes | Bacteroidia | Flavobacteriales | Flavobacteriaceae | NA |
| ASV_766 | Low | 0.001 | Actinobacteria | Acidimicrobiia | Microtrichales | Microtrichaceae | Sva0996_marine_group |
| ASV_772 | Low | 0.001 | Planctomycetes | Planctomycetacia | Pirellulales | Pirellulaceae | Blastopirellula |
| ASV_773 | High | 0.006 | Proteobacteria | Alphaproteobacteria | Sphingomonadales | Sphingomonadaceae | NA |
| ASV_774 | Low | 0.001 | Bacteroidetes | Bacteroidia | Chitinophagales | Saprospiraceae | Lewinella |
| ASV_777 | Low | 0.001 | Planctomycetes | Planctomycetacia | Planctomycetales | NA | NA |
| ASV_779 | Low | 0.001 | Proteobacteria | Alphaproteobacteria | Rhodobacterales | Rhodobacteraceae | Roseobacter_clade_NAC 11-7_lineage |
| ASV_783 | Low | 0.016 | Bacteroidetes | Bacteroidia | Flavobacteriales | Flavobacteriaceae | Jejudonia |
| ASV_786 | Low | 0.001 | Proteobacteria | Alphaproteobacteria | Rhodobacterales | Rhodobacteraceae | NA |
| ASV_787 | High | 0.035 | Proteobacteria | Gammaproteobacteria | Oceanospirillales | Alcanivoracaceae | Alcanivorax |
| ASV_796 | Low | 0.004 | Bacteroidetes | Bacteroidia | Flavobacteriales | Flavobacteriaceae | Aquibacter |
| ASV_799 | Low | 0.001 | Proteobacteria | Gammaproteobacteria | Gammaproteobacteria_ Incertae_Sedis | Unknown_Family | Candidatus_Berkiella |
| ASV_802 | Middle | 0.025 | Bacteroidetes | Bacteroidia | Flavobacteriales | Flavobacteriaceae | Nonlabens |
| ASV_809 | Low | 0.002 | Bacteroidetes | Bacteroidia | Flavobacteriales | Flavobacteriaceae | NA |
| ASV_813 | Middle | 0.032 | Bacteroidetes | Bacteroidia | Chitinophagales | Saprospiraceae | Lewinella |
| ASV_820 | Low | 0.001 | Proteobacteria | Alphaproteobacteria | Rhodobacterales | Rhodobacteraceae | Lentibacter |
| ASV_823 | Low | 0.002 | Bacteroidetes | Bacteroidia | Flavobacteriales | Flavobacteriaceae | Marixanthomonas |
| ASV_824 | High | 0.048 | Proteobacteria | Gammaproteobacteria | Alteromonadales | Alteromonadaceae | Salinimonas |
| ASV_826 | Middle | 0.001 | Proteobacteria | Alphaproteobacteria | Caulobacterales | Hyphomonadaceae | Litorimonas |
| ASV_829 | Low | 0.001 | Bacteroidetes | Bacteroidia | Chitinophagales | Saprospiraceae | Rubidimonas |
| ASV_831 | Middle | 0.002 | Proteobacteria | Gammaproteobacteria | Gammaproteobacteria_ Incertae_Sedis | Unknown_Family | Marinicella |
| ASV_836 | Middle | 0.001 | Bacteroidetes | Bacteroidia | Flavobacteriales | Flavobacteriaceae | Dokdonia |
| ASV_841 | Middle | 0.046 | Cyanobacteria | Oxyphotobacteria | Phormidesmiales | Phormidesmiaceae | Phormidesmis_ANT.LA CV5.1 |
| ASV_845 | Low | 0.001 | Proteobacteria | Alphaproteobacteria | Rhodobacterales | Rhodobacteraceae | NA |
| ASV_850 | Middle | 0.002 | Proteobacteria | Alphaproteobacteria | Rhizobiales | Rhizobiaceae | Pseudahrensia |

| | | | | | | | |
|---|---|---|---|---|---|---|---|
| ASV_857 | Low | 0.001 | Verrucomicrobia | Verrucomicrobiae | Verrucomicrobiales | Rubritaleaceae | Rubritalea |
| ASV_859 | Middle | 0.013 | Bacteroidetes | Bacteroidia | Chitinophagales | Saprospiraceae | Lewinella |
| ASV_860 | Middle | 0.015 | Proteobacteria | Alphaproteobacteria | NA | NA | NA |
| ASV_861 | Low | 0.001 | Bacteroidetes | Bacteroidia | Flavobacteriales | Flavobacteriaceae | Aquimarina |
| ASV_866 | High | 0.005 | Cyanobacteria | Oxyphotobacteria | Nostocales | Xenococcaceae | Pleurocapsa_PCC-7319 |
| ASV_867 | Low | 0.001 | Deinococcus-Thermus | Deinococci | Deinococcales | Trueperaceae | Truepera |
| ASV_868 | Middle | 0.02 | Proteobacteria | Alphaproteobacteria | Rhizobiales | Rhizobiaceae | Pseudahrensia |
| ASV_869 | Middle | 0.005 | Proteobacteria | Alphaproteobacteria | Rhodobacterales | Rhodobacteraceae | Roseobacter |
| ASV_873 | Low | 0.01 | Proteobacteria | Gammaproteobacteria | Thiotrichales | Thiotrichaceae | Cocleimonas |
| ASV_874 | Low | 0.001 | Planctomycetes | Planctomycetacia | Pirellulales | Pirellulaceae | Rhodopirellula |
| ASV_878 | Low | 0.001 | Proteobacteria | Alphaproteobacteria | NA | NA | NA |
| ASV_893 | Low | 0.011 | Proteobacteria | Alphaproteobacteria | Rhodobacterales | Rhodobacteraceae | Actibacterium |
| ASV_902 | Low | 0.014 | Bacteroidetes | Bacteroidia | Flavobacteriales | Flavobacteriaceae | NA |
| ASV_906 | Middle | 0.001 | Proteobacteria | Alphaproteobacteria | Caulobacterales | Hyphomonadaceae | Litorimonas |
| ASV_909 | Low | 0.002 | Proteobacteria | Alphaproteobacteria | Caulobacterales | Hyphomonadaceae | NA |
| ASV_912 | Low | 0.001 | Planctomycetes | Planctomycetacia | Pirellulales | Pirellulaceae | Blastopirellula |
| ASV_914 | Low | 0.005 | Bacteroidetes | Bacteroidia | Flavobacteriales | Flavobacteriaceae | Ulvibacter |
| ASV_918 | Low | 0.003 | Proteobacteria | Alphaproteobacteria | Rhodobacterales | Rhodobacteraceae | Loktanella |
| ASV_919 | Low | 0.001 | Proteobacteria | Alphaproteobacteria | Rhizobiales | Rhizobiaceae | Pseudahrensia |
| ASV_921 | High | 0.024 | Planctomycetes | NA | NA | NA | NA |
| ASV_923 | Middle | 0.009 | Bacteroidetes | Bacteroidia | Flavobacteriales | Flavobacteriaceae | NA |
| ASV_926 | Middle | 0.011 | Proteobacteria | Alphaproteobacteria | Rhizobiales | Rhizobiaceae | Pseudahrensia |
| ASV_927 | Low | 0.001 | Bacteroidetes | Bacteroidia | Chitinophagales | Saprospiraceae | NA |
| ASV_928 | High | 0.024 | Proteobacteria | Alphaproteobacteria | Sphingomonadales | Sphingomonadaceae | NA |
| ASV_929 | Low | 0.002 | Proteobacteria | Gammaproteobacteria | Thiohalorhabdales | Thiohalorhabdaceae | Granulosicoccus |
| ASV_935 | High | 0.015 | Cyanobacteria | Oxyphotobacteria | Nostocales | Xenococcaceae | Pleurocapsa_PCC-7319 |
| ASV_936 | High | 0.015 | Bacteroidetes | Bacteroidia | Flavobacteriales | Flavobacteriaceae | Pricia |
| ASV_937 | Low | 0.003 | Proteobacteria | Alphaproteobacteria | Sphingomonadales | Sphingomonadaceae | Altererythrobacter |
| ASV_943 | Low | 0.022 | Bacteroidetes | Bacteroidia | Flavobacteriales | Flavobacteriaceae | Psychroserpens |
| ASV_944 | Middle | 0.041 | Bacteroidetes | Bacteroidia | Flavobacteriales | Flavobacteriaceae | Dokdonia |
| ASV_946 | Middle | 0.014 | Bacteroidetes | Bacteroidia | Chitinophagales | Saprospiraceae | Lewinella |
| ASV_947 | High | 0.032 | Proteobacteria | Alphaproteobacteria | Rhodobacterales | Rhodobacteraceae | Jannaschia |
| ASV_948 | Low | 0.001 | Bacteroidetes | Bacteroidia | Chitinophagales | Saprospiraceae | NA |
| ASV_955 | Low | 0.001 | Bacteroidetes | Bacteroidia | Flavobacteriales | Flavobacteriaceae | Marixanthomonas |
| ASV_962 | Low | 0.001 | Planctomycetes | Planctomycetacia | Pirellulales | Pirellulaceae | Blastopirellula |
| ASV_970 | Low | 0.003 | Proteobacteria | Alphaproteobacteria | Rhizobiales | Rhizobiaceae | Pseudahrensia |
| ASV_974 | High | 0.035 | Proteobacteria | Gammaproteobacteria | NA | NA | NA |
| ASV_976 | Low | 0.001 | Proteobacteria | Gammaproteobacteria | Thiotrichales | Thiotrichaceae | Leucothrix |
| ASV_977 | Middle | 0.014 | Proteobacteria | Alphaproteobacteria | Sphingomonadales | Sphingomonadaceae | Altererythrobacter |

| | | | | | | | |
|---|---|---|---|---|---|---|---|
| ASV_980 | Low | 0.008 | Proteobacteria | Gammaproteobacteria | KI89A_clade | NA | NA |
| ASV_990 | Low | 0.001 | Bacteroidetes | Bacteroidia | Flavobacteriales | Flavobacteriaceae | Ulvibacter |
| ASV_991 | Low | 0.001 | Bacteroidetes | Bacteroidia | Flavobacteriales | Flavobacteriaceae | Pibocella |
| ASV_993 | Low | 0.001 | Planctomycetes | Planctomycetacia | Pirellulales | Pirellulaceae | Blastopirellula |
| ASV_995 | Middle | 0.005 | Cyanobacteria | Oxyphotobacteria | Phormidesmiales | Phormidesmiaceae | Phormidesmis_ANT.LA CV5.1 |
| ASV_1000 | Middle | 0.016 | Bacteroidetes | Bacteroidia | Flavobacteriales | Flavobacteriaceae | Maribacter |
| ASV_1007 | Low | 0.038 | Bacteroidetes | Bacteroidia | Flavobacteriales | Flavobacteriaceae | Aquibacter |
| ASV_1018 | Low | 0.003 | Bacteroidetes | Bacteroidia | Flavobacteriales | Flavobacteriaceae | Aquibacter |
| ASV_1028 | Low | 0.002 | Bacteroidetes | Bacteroidia | Flavobacteriales | Flavobacteriaceae | Maritimimonas |
| ASV_1037 | Middle | 0.002 | Verrucomicrobia | Verrucomicrobiae | Verrucomicrobiales | Rubritaleaceae | Haloferula |
| ASV_1048 | Low | 0.001 | Proteobacteria | Alphaproteobacteria | Rhodobacterales | Rhodobacteraceae | NA |
| ASV_1053 | High | 0.002 | Cyanobacteria | Oxyphotobacteria | Nostocales | Xenococcaceae | Pleurocapsa_PCC-7319 |
| ASV_1064 | Middle | 0.016 | Bacteroidetes | Bacteroidia | Flavobacteriales | Flavobacteriaceae | Jejudonia |
| ASV_1067 | Low | 0.004 | Bacteroidetes | Bacteroidia | Flavobacteriales | Flavobacteriaceae | Psychroserpens |
| ASV_1081 | Low | 0.001 | Proteobacteria | Gammaproteobacteria | NA | NA | NA |
| ASV_1082 | High | 0.039 | Proteobacteria | Alphaproteobacteria | Rhodobacterales | Rhodobacteraceae | NA |
| ASV_1083 | Low | 0.001 | Proteobacteria | Alphaproteobacteria | Rhodobacterales | Rhodobacteraceae | Amylibacter |
| ASV_1084 | Low | 0.003 | Bacteroidetes | Bacteroidia | Chitinophagales | Saprospiraceae | NA |
| ASV_1089 | Low | 0.001 | Proteobacteria | Gammaproteobacteria | NA | NA | NA |
| ASV_1093 | Middle | 0.017 | Proteobacteria | Gammaproteobacteria | NA | NA | NA |
| ASV_1098 | Low | 0.006 | Bacteroidetes | Bacteroidia | Flavobacteriales | Flavobacteriaceae | Lutibacter |
| ASV_1106 | Low | 0.001 | Acidobacteria | Thermoanaerobaculia | Thermoanaerobaculale s | Thermoanaerobaculac eae | Subgroup_10 |
| ASV_1111 | Low | 0.002 | Planctomycetes | Planctomycetacia | Pirellulales | Pirellulaceae | Blastopirellula |
| ASV_1112 | Low | 0.002 | Proteobacteria | Gammaproteobacteria | Thiotrichales | Thiotrichaceae | Cocleimonas |
| ASV_1116 | Middle | 0.007 | Proteobacteria | Alphaproteobacteria | Rhizobiales | Rhizobiaceae | Pseudahrensia |
| ASV_1135 | Low | 0.001 | Proteobacteria | Gammaproteobacteria | Thiotrichales | Thiotrichaceae | Cocleimonas |
| ASV_1148 | Low | 0.001 | Proteobacteria | Alphaproteobacteria | Rhodobacterales | Rhodobacteraceae | Lentibacter |
| ASV_1164 | Low | 0.004 | Proteobacteria | Gammaproteobacteria | Thiotrichales | Thiotrichaceae | Thiothrix |
| ASV_1188 | Middle | 0.001 | Bacteroidetes | Bacteroidia | Flavobacteriales | Flavobacteriaceae | Kordia |
| ASV_1189 | Low | 0.001 | Proteobacteria | Alphaproteobacteria | NA | NA | NA |
| ASV_1194 | Low | 0.001 | Proteobacteria | Alphaproteobacteria | Sphingomonadales | Sphingomonadaceae | Altererythrobacter |
| ASV_1207 | Low | 0.046 | Bacteroidetes | Bacteroidia | Chitinophagales | Saprospiraceae | Lewinella |
| ASV_1234 | Middle | 0.003 | Proteobacteria | Gammaproteobacteria | Alteromonadales | Alteromonadaceae | Glaciecola |
| ASV_1238 | Middle | 0.002 | Proteobacteria | Alphaproteobacteria | Rhodobacterales | Rhodobacteraceae | NA |
| ASV_1239 | Middle | 0.047 | Proteobacteria | Gammaproteobacteria | Cellvibrionales | Spongiibacteraceae | NA |
| ASV_1242 | Low | 0.001 | Planctomycetes | Planctomycetacia | Pirellulales | Pirellulaceae | Blastopirellula |
| ASV_1246 | Low | 0.005 | Proteobacteria | Alphaproteobacteria | Rhodobacterales | Rhodobacteraceae | Octadecabacter |
| ASV_1263 | Low | 0.001 | Cyanobacteria | Oxyphotobacteria | Synechococcales | Synechococcales_Ince rtae_Sedis | Schizothrix_LEGE_0716 4 |
| ASV_1275 | Low | 0.009 | Proteobacteria | Gammaproteobacteria | KI89A_clade | NA | NA |

| | | | | | | | |
|---|---|---|---|---|---|---|---|
| ASV_1343 | Low | 0.013 | Planctomycetes | Planctomycetacia | Planctomycetales | NA | NA |
| ASV_1347 | Low | 0.002 | Bacteroidetes | Bacteroidia | Chitinophagales | Saprospiraceae | Rubidimonas |
| ASV_1348 | Low | 0.001 | Proteobacteria | Alphaproteobacteria | Rhodobacterales | Rhodobacteraceae | NA |
| ASV_1354 | Low | 0.017 | Planctomycetes | OM190 | NA | NA | NA |
| ASV_1395 | Low | 0.001 | Proteobacteria | Alphaproteobacteria | NA | NA | NA |
| ASV_1420 | High | 0.048 | Proteobacteria | Alphaproteobacteria | Micavibrionales | NA | NA |
| ASV_1456 | Low | 0.009 | Proteobacteria | Gammaproteobacteria | Cellvibrionales | Spongiibacteraceae | NA |
| ASV_1498 | Middle | 0.005 | Proteobacteria | Alphaproteobacteria | Rhodobacterales | Rhodobacteraceae | Roseobacter |
| ASV_1612 | Middle | 0.024 | Bacteroidetes | Bacteroidia | Flavobacteriales | Flavobacteriaceae | Dokdonia |
| ASV_1846 | Low | 0.028 | Planctomycetes | Planctomycetacia | Pirellulales | Pirellulaceae | Blastopirellula |
| ASV_1915 | Low | 0.022 | Proteobacteria | Alphaproteobacteria | NA | NA | NA |
| ASV_2088 | Low | 0.006 | Planctomycetes | Planctomycetacia | Pirellulales | Pirellulaceae | Blastopirellula |

### Appendix S4

**Figure S1.** Degree distribution of the co-occurrence network of A) macroorganisms and B) microorganisms.

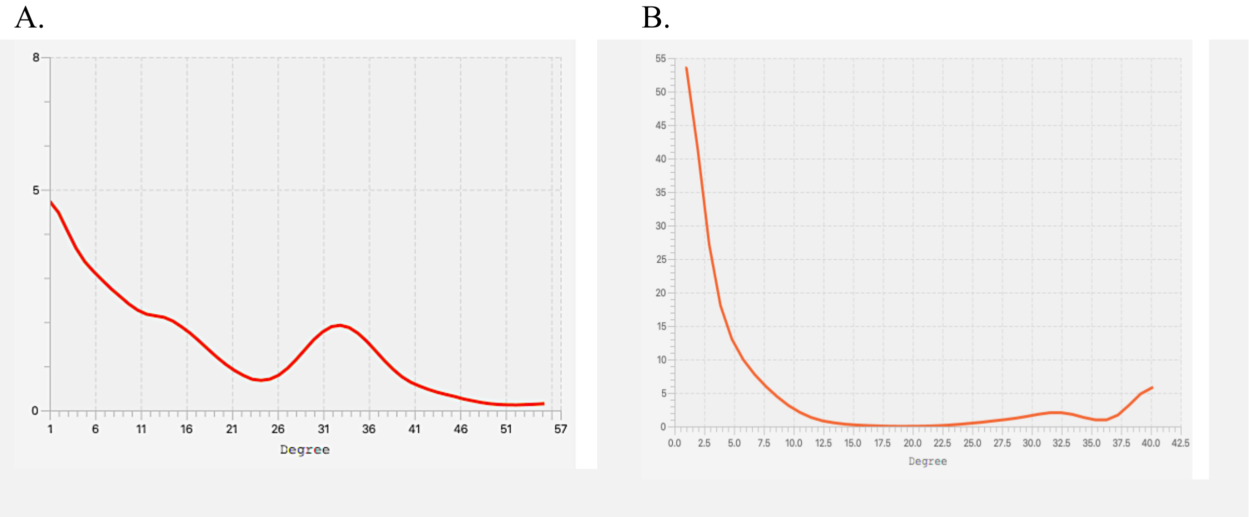

**Appendix S4**

**Figure S2.** Percent of species/OTUs in each cluster derived using the leading eigenvalue method at each tidal height that are environmental specialists at each tidal height and that are non-specialists. Both networks are modular with A) the macroorganisms network weighted by correlation strength forming 6 clusters and B) the microorganism network weighted by correlation strength forming 3 clusters. These clusters show some relation to tidal height groupings, but they are not significantly related to tidal height groups. H: High intertidal zone specialist, M: Middle intertidal zone specialist, L: Low intertidal zone specialist, N: Not specialist.

**A)** **B)**

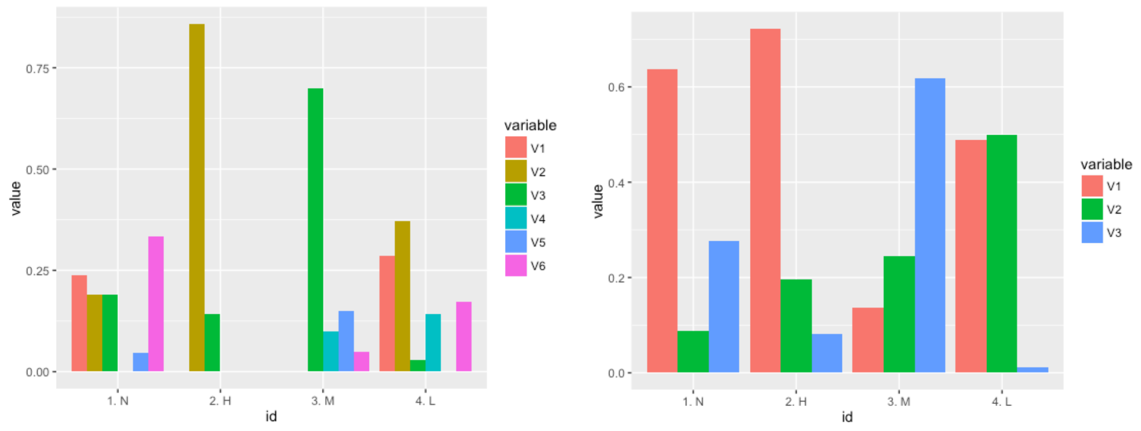
